# K2P Channels Regulate Presynaptic Organisation through a Membrane Potential-Independent Mechanism

**DOI:** 10.64898/2026.08.21.746268

**Authors:** Jun Meng, Neeraja Ramakrishnan, Yi Li, Thomas Boulin, Shangbang Gao, Mei Zhen, Isabel Beets, William R Schafer

## Abstract

Neuronal ion channels have well-established effects on synaptic plasticity, in many cases by influencing pathways that depend on membrane excitability. Here we find that a *C. elegans* two-pore domain potassium (K2P) channel, TWK-40, regulates presynaptic organisation through a membrane potential-independent mechanism. Instead, this mechanism depends on TWK-40’s effects on intracellular potassium levels. Loss-of-function mutations in *twk-40* lead to excessive presynaptic protein accumulation, while gain-of-function mutations lead to depleted presynaptic components and cause synaptic transmission deficits. These changes in presynaptic protein abundance are phenocopied by transporter mutations predicted to alter intracellular potassium homeostasis in the same direction as TWK-40 mutations, but not by sodium channel mutations that mimic its effects on membrane excitability. This indicates that cytoplasmic potassium promotes presynaptic assembly. This process depends on pyruvate kinase PYK-1, a potassium-sensitive enzyme, and three transcription factors. These findings establish a new pathway linking neuronal potassium homeostasis to the control of presynaptic organisation and synaptic function.

## Introduction

The assembly of neural circuits involves mechanisms that are both activity dependent and activity independent^1^. Activity-dependent programs have been more extensively characterised^1–4^. For example, neuronal activity induces calcium-dependent gene expression that reshapes synaptic connectivity through transcription factors such as CREB^5^, MEF2^6^, and cytoskeletal effectors such as Arc^7^. In parallel, activity-independent cues have been shown to influence synaptic development. Specifically, synaptic target selection and initial synaptic differentiation take place in the absence of electrical activity^8–10^. These activity-independent processes have been proposed to establish early wiring patterns that are later refined by neuronal activity, but mechanisms mediating these processes are not understood.

Ion channels are best known for their rapid regulation of neuronal excitability, but whether they also exert persistent effects on neuronal architecture remains less clear. Background ion channels control resting membrane potential, while also contributing to the homeostasis of cytoplasmic ions^11^. Among ion channels, members of the two-pore domain potassium (K2P) channel family are major contributors to background potassium conductance and thus play key roles in establishing the steady-state membrane potential^12–16^. K2P channels share evolutionarily conserved sequence homology across species and are broadly expressed across the nervous system^15–17^. Structurally, K2P channels assemble as dimers, with each subunit containing two pore-forming domains. Some K2P channels exhibit functionally specialised subcellular localisation, such as to nodes of Ranvier to support action potential propagation^18^, suggesting compartment-specific physiological functions. However, the possibility that K2P channels play direct roles in the spatial organisation of synaptic architecture remains largely unexplored.

To explore this question, we turned to *Caenorhabditis elegans*, a genetically tractable animal with a small and well-characterised nervous system^19^. The *C. elegans* genome encodes ∼ 47 K2P-encoding genes, a striking expansion of this gene family compared with only 15 in humans and 11 in *Drosophila*^20,21^, despite its nervous system comprising just a few hundred neurons^22^. Most of these genes remain uncharacterised, and loss-of-function mutations often result in subtle or no phenotype, possibly due to functional redundancy^20^. One of the few exceptions is TWK-40, a K2P channel expressed primarily in the motor circuits^23–25^. TWK-40 has been shown to specifically function in the AVA premotor interneurons, where it contributes to the maintenance of its membrane voltage within a narrow range^23,24^. The AVA neurons form part of the highly connected rich-club core of the synaptic connectome and are also among the most highly connected neurons in the neuropeptidergic network^22,26,27^. Loss-and gain-of-function alleles of the *twk-40* gene yield opposite effects on backward locomotion^23,24^, providing a genetically tractable model to investigate how K2P channel activity influences synaptic organisation.

Here, we show that TWK-40 is enriched at presynaptic regions along the ventral nerve cord and regulates presynaptic organisation in the AVA premotor interneurons. Unexpectedly, we find that this regulation occurs independently of changes in membrane potential and instead depends on TWK-40’s control of the intracellular potassium concentration. The potassium-sensitive enzyme PYK-1 is required for these effects, suggesting a possible mechanistic link between potassium homeostasis and synaptic organisation. Altered TWK-40 activity, associated with disrupted K⁺ homeostasis, alters the expression of three transcription factors that affect synapse development in AVA neurons. These findings uncover a membrane potential-independent, *in vivo* mechanism by which a K2P channel regulates chemical synapse development and function.

## Results

### Increased TWK-40 Activity Inhibits Chemical Synaptic Function

We previously showed that the *twk-40* loss-of-function allele *hp834* [hereafter *twk-40(lf)*] and the gain-of-function allele *bln336*, which carries the L159N substitution [hereafter *twk-40(gf, L159N)*], displayed opposite locomotor phenotypes: *twk-40(lf)* animals showed an increased frequency of spontaneous backward movement, whereas *twk-40(gf, L159N)* animals failed to initiate backward movement^23^ (Figure S1A). The AVA premotor interneurons, whose membrane potential is altered by mutations in *twk-40*, make both electrical and chemical synapses onto the motor neurons that execute backward locomotion, and these chemical synapses in particular are critical for initiating a reversal^28,29^. For example, *C. elegans* exhibits a robust head-touch-evoked escape response that requires AVA activation^30^, and *twk-40(gf, L159N)* mutants are strongly defective in this reversal response (Figure S1B). Because of the severity of this phenotype, and the fact that neuronal activity is known to influence synapse development and function^4,31^, these results suggested that excess TWK-40 activity might disrupt AVA’s ability to drive chemical synaptic output.

To investigate this possibility, we examined the effect of optogenetic depolarisation of AVA using Chrimson, a red-shifted channelrhodopsin^32^. When AVA-Chrimson-expressing worms were stimulated by light, they displayed robust backward locomotion, similar to the reversal evoked by head touch (Figure 1A, Figure S1B–D). However, in AVA-TWK-40(gf, L159N) animals, Chrimson-mediated AVA activation failed to elicit backward movement (Figure 1A, Figure S1D), despite evoking clear calcium transients in AVA (Figure 1B; Figure S1E, F). Although these calcium responses were reduced relative to wild type, their presence indicates that the behavioural failure is not due to a complete lack of AVA activation. These results suggest that impaired backward movement cannot be explained solely by reduced AVA activation and may also involve defects in synaptic transmission.

**Figure 1.**
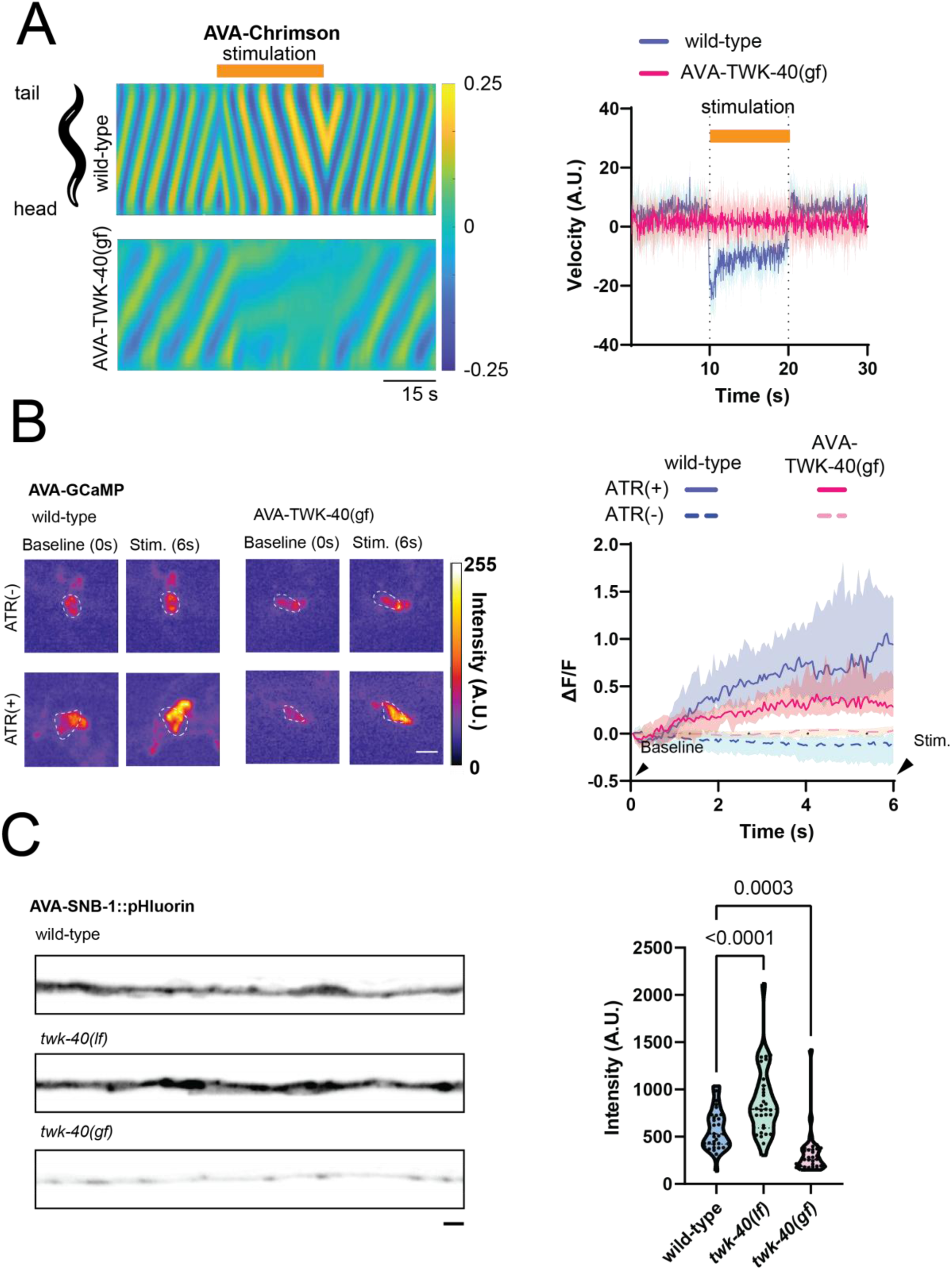
TWK-40 activity regulates neural function. (A) **Left:** Kymographs displaying the curvature map of animals expressing Chrimson in AVA for wild-type and AVA-specific TWK-40(gf) genotypes. In wild-type animals, AVA stimulation induces backward movement, whereas in AVA-TWK-40(gf) animals, AVA stimulation fails to elicit backward movement. **Right:** Velocity traces showing the median and 95% confidence interval (CI) for each genotype during AVA stimulation. **Sample sizes:** wild-type (N = 10 animals), AVA-TWK-40(gf) (N = 11 animals). (B) **Left:** Representative images showing AVA calcium activity (GCaMP) at the beginning and during continuous photostimulation in animals supplemented with ATR (+) or without ATR (−). The illumination used for calcium imaging simultaneously activates Chrimson and is maintained throughout the recording. Baseline indicates the first frame at stimulation onset; Stim. indicates a later time point during Chrimson stimulation. All-trans-retinal (ATR) is a cofactor of Chrimson. Without ATR, calcium levels remain unchanged, whereas with ATR, calcium levels increase upon AVA stimulation. The circles denote the AVA neurons; the adjacent uncircled signal corresponds to AVE. Scale bar: 5 μm. **Right:** calcium traces (ΔF/F) over time with median and 95% CI. Wild-type is shown in blue and AVA-TWK-40(gf) in magenta; solid and dashed lines indicate ATR(+) and ATR(−), respectively. Shaded areas indicate the 95% CI. AVA stimulation in the presence of ATR leads to an increase in calcium levels in both wild-type and AVA-TWK-40(gf) animals. However, in AVA-TWK-40(gf) animals, the increase is attenuated compared to wild-type. **Sample sizes:** ATR(-): wild-type (N = 22 animals), AVA-TWK-40(gf) (N = 25 animals). ATR(+): wild-type (N = 33 animals), AVA-TWK-40(gf) (N = 30 animals). (C) **Left:** Representative confocal images showing maximum intensity projections of SNB-1::pHluorin in AVA along the ventral nerve cord for wild-type, twk-40(lf), and twk-40(gf) genotypes. Scale bar: 2 μm. **Right:** Quantification of SNB-1::pHluorin intensity, showing significantly increased SNB-1::pHluorin fluorescence in twk-40(lf) and decreased fluorescence in twk-40(gf) compared with wild type. **Sample sizes:** wild-type (N = 35 animals), twk-40(lf) (N = 31 animals), twk-40(gf) (N = 32 animals).

To directly assess the effect of *twk-40* on chemical synaptic output, we examined synaptic vesicle exocytosis using SNB-1::pHluorin, a fusion of the synaptic vesicle protein synaptobrevin/SNB-1 to the pH-sensitive GFP variant superecliptic pHluorin, whose fluorescence increases upon vesicle fusion and exposure to extracellular pH^33^. We observed that signal intensity in the ventral nerve cord, corresponding to synaptic vesicle release specifically from AVA, was significantly decreased in *twk-40(gf, L159N)* mutants and increased in *twk-40(lf)* animals (Figure 1C). These findings reveal TWK-40-dependent changes in the presynaptic SNB-1::pHluorin signal, consistent with altered synaptic vesicle exocytosis and/or vesicle abundance^28,29^. Although reduced AVA excitability likely contributes to the backward-locomotion defect in *twk-40(gf, L159N)* mutants, the failure of direct optogenetic AVA activation to restore backward movement suggests an additional defect downstream of AVA activation, potentially involving presynaptic output or chemical synaptic transmission.

### TWK-40 Activity Regulates Presynaptic Organisation

Some K2P channels exhibit subcellular compartmentalisation that shapes their physiological roles^34^. To determine whether TWK-40 is positioned to influence synaptic function, we examined its distribution along the ventral nerve cord, where AVA and other *twk-40*-expressing neurons form presynaptic termini. Using an endogenously tagged TWK-40::tagRFP reporter^23,35^, and SNB-1::GFP expressed under the *twk-40* promoter, we observed overlapping puncta along the ventral nerve cord by immunostaining (Figure 2A). These results are consistent with TWK-40’s enrichment at presynaptic regions and a potential role in presynaptic organisation.

**Figure 2.**
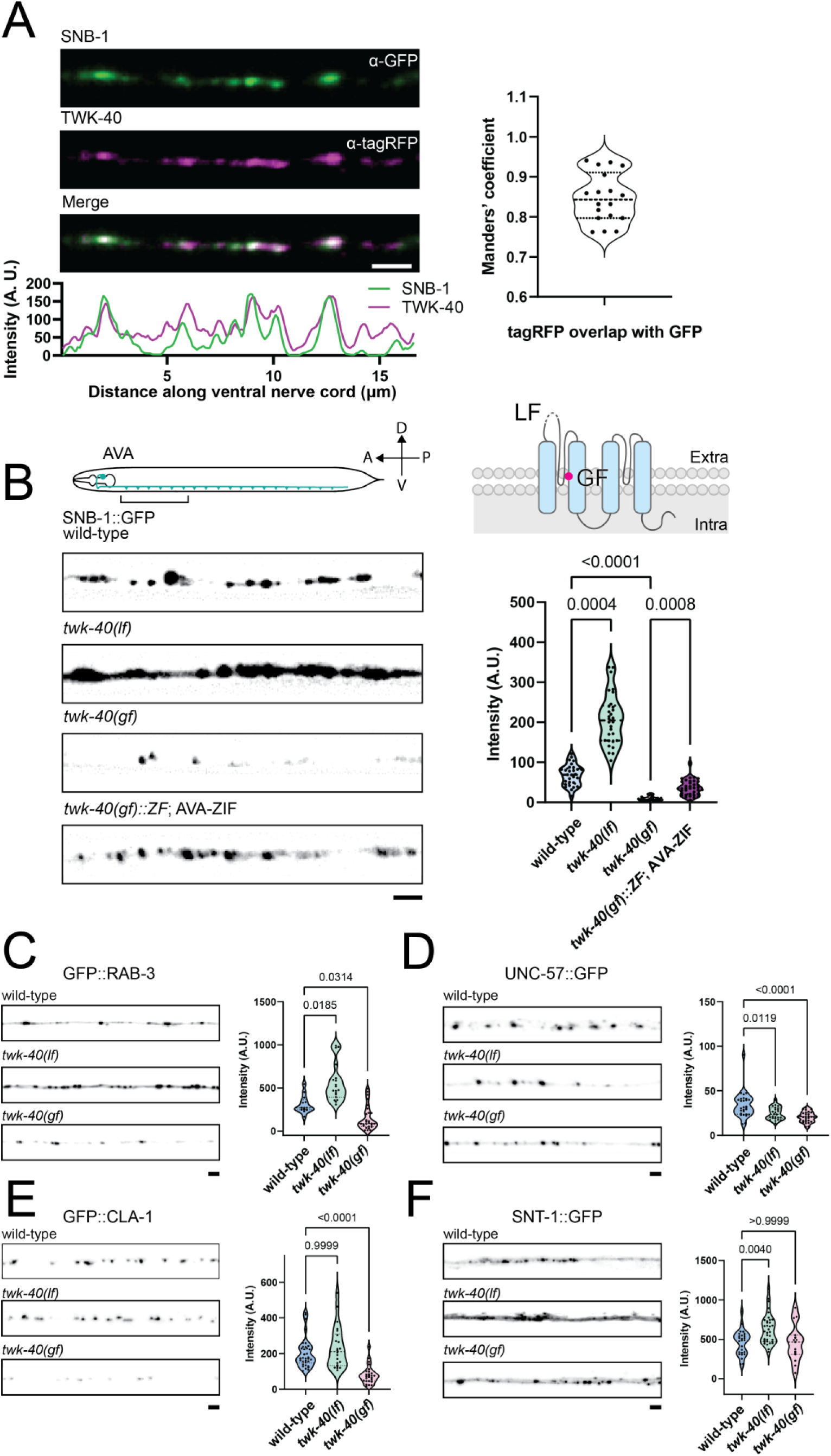
The K2P channel TWK-40 localises to the presynapse and regulates the presynaptic organisation. (A) **Left:** Representative confocal immunostaining images showing SNB-1::GFP (green) and TWK-40::tagRFP::ZF (magenta) localisation along the ventral nerve cord in the twk-40::tagRFP::ZF; Ptwk-40-SNB-1::GFP genotype. **Top:** SNB-1::GFP signal detected using α-GFP. **Middle:** TWK-40::tagRFP signal detected using α-tagRFP. **Bottom:** Merged image illustrating colocalisation of the two signals. Scale bar: 2 μm. **Right:** Quantification of colocalisation between TWK-40 and SNB-1, measured by Manders’ coefficient. N = 18 animals. (B) **Top left:** Schematic illustration of SNB-1 expressed in AVA neurons and the corresponding ventral nerve cord region where confocal images were taken and analysed. **Top right:** Schematic representation of TWK-40 loss-of-function (lf) and gain-of-function (gf) mutations. **Bottom left:** Representative confocal images of AVA-specific SNB-1::GFP signal along the ventral nerve cord in each genetic background: wild-type, twk-40(lf), twk-40(gf), and twk-40(gf)::ZF; AVA-ZIF (degraded). Scale bar: 2 μm. **Bottom right:** Quantification of SNB-1::GFP intensity for each genotype. The SNB-1::GFP signal intensity in AVA neurons is significantly increased in twk-40(lf) mutants and decreased in twk-40(gf) mutants. Specific degradation of TWK-40(gf) in AVA neurons restores SNB-1::GFP intensity towards wild-type levels. **Sample sizes:** wild-type (N = 29 animals), twk-40(lf) (N = 30 animals), twk-40(gf) (N = 10 animals), twk-40(gf)::ZF; AVA-ZIF (degraded) (N = 37 animals). (C–F) **Left:** Representative confocal images of presynaptic markers along the ventral nerve cord in AVA neurons from wild-type, twk-40(lf), and twk-40(gf) backgrounds. Markers include GFP::RAB-3 (C), UNC-57::GFP (D), GFP::CLA-1 (E), and SNT-1::GFP (F). Scale bar: 2 μm. **Right:** Quantification of fluorescence intensity for each presynaptic marker, demonstrating significant differences between wild-type and twk-40 mutants. **Sample sizes:** GFP::RAB-3: wild-type (N = 13 animals), twk-40(lf) (N = 24 animals), twk-40(gf) (N = 27 animals). UNC-57::GFP: wild-type (N = 28 animals), twk-40(lf) (N = 26 animals), twk-40(gf) (N = 31 animals). GFP::CLA-1: wild-type (N = 34 animals), twk-40(lf) (N = 22 animals), twk-40(gf) (N = 26 animals). SNT-1::GFP: wild-type (N = 27 animals), twk-40(lf) (N = 35 animals), twk-40(gf) (N = 15 animals).

We next assessed presynaptic organisation, as reflected by the distribution and accumulation of presynaptic protein markers along the AVA ventral nerve cord. We quantified SNB-1::GFP in AVA across *twk-40* loss-and gain-of-function mutants. We observed that SNB-1::GFP fluorescence was significantly elevated in *twk-40(lf)* animals and markedly reduced in *twk-40(gf, L159N)* animals (Figure 2B). Changes in puncta density and size were also observed (Figure S2B). To exclude the possibility that the decrease in SNB-1 fluorescent signal reflected GFP quenching rather than a true loss of presynaptic protein, we performed anti-GFP immunostaining, which recapitulated the reduction observed by native fluorescence (Figure S2A). These results confirmed a bona fide decrease in presynaptic SNB-1 levels in *twk-40(gf, L159N)* animals.

We next asked whether this reduction was due to excessive TWK-40 activity specifically within AVA. We used the ZF/ZIF-1 degradation system to selectively degrade TWK-40(gf, L159N) in AVA and assayed the effect of this depletion on SNB-1::GFP levels. We observed a robust rescue of the *twk-40(gf, L159N)* phenotype, with SNB-1::GFP intensity markedly increased in TWK-40(gf, L159N) AVA-depleted animals compared to *twk-40(gf, L159N)* (Figure 2B). These results indicate that TWK-40 regulates presynaptic SNB-1 levels in a cell-autonomous manner.

To assess whether the effect of TWK-40 is unique to SNB-1 or indicative of a broader role in presynaptic organisation, we examined additional presynaptic markers, including RAB-3 (RAB3A homolog^36,37^), UNC-57 (endophilin A^38^), SNT-1 (synaptotagmin 1^39^), and CLA-1 (active zone protein^40^). These markers showed marker-specific changes, with RAB-3 most closely recapitulating the SNB-1 phenotype (Figure 2C–F, Figure S2C–F). Given the robust phenotype and the central role of SNB-1 in synaptic vesicle exocytosis, we focused on SNB-1::GFP as the primary readout for presynaptic structure in subsequent experiments.

### TWK-40 Regulates Synaptic Organisation and Function During Larval Development

The AVA neurons are born embryonically and form functional synapses in the newly hatched L1 larva. We therefore examined whether TWK-40 acts during a specific developmental stage to regulate presynaptic structure and function. We used the ZF/ZIF-1 degradation system coupled with a heat shock promoter to deplete TWK-40(gf, L159N) at different developmental stages (Figure 3A), and examined SNB-1::GFP levels and locomotor behaviour at the final larval stage (L4). When heat shock was applied at later stages (L3 and L4), SNB-1::GFP expression partially recovered (Figure 3C), and locomotion improved (Figure 3B). Thus, heat-shock-induced depletion of TWK-40(gf) at later larval stages partially ameliorated the presynaptic defects. Conversely, animals heat-shocked at L1 or L2 and subsequently maintained at 20°C until L4 showed reduced SNB-1::GFP levels and abnormal locomotion. Together, these results indicate that TWK-40 activity continues to influence presynaptic organisation and locomotor function during later larval development.

**Figure 3.**
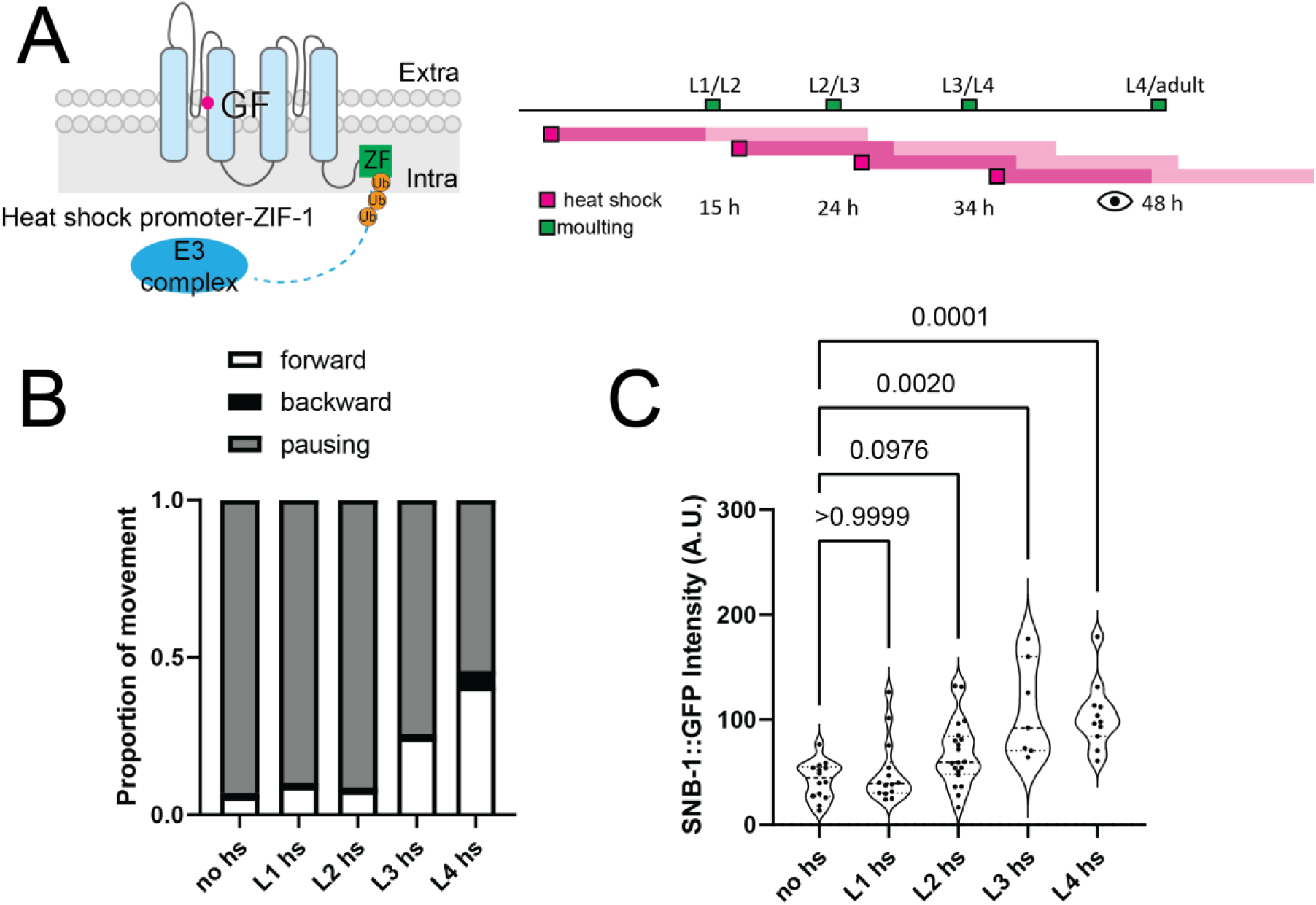
TWK-40 regulates presynaptic organisation and locomotor function during larval development. (A) **Left:** Schematic diagram illustrating the heat shock-induced ZF/ZIF-1 degradation system, which enables temporal control of TWK-40(gf) levels. The heat shock promoter drives ZIF-1 expression, which, in combination with an E3 ubiquitin ligase complex, targets TWK-40(gf)::ZF for degradation. **Right:** Timeline showing developmental stages (L1 to adult) with magenta boxes indicating heat shock application time points. Light magenta shading schematically indicates the approximate period during which ZF1-mediated TWK-40 depletion is expected following heat-shock-induced ZIF-1 expression, allowing TWK-40(gf) levels to be modulated at specific developmental stages. (B) Proportion of movement (forward, backward, pausing) for animals with the genotype twk-40(gf)::ZF; Phsp-ZIF-1 under different heat shock conditions, pooled across 10 animals per condition. Heat shock application at the L3 or L4 stage partially rescues the locomotor phenotype, increasing the proportion of forward movement and reducing pausing. **Sample sizes:** N = 10 animals per condition. (C) Quantification of AVA-SNB-1::GFP intensity under different heat shock conditions. Heat shock at the L3 and L4 stages partially and significantly rescues the presynaptic phenotype. **Sample sizes:** No heat shock control (N = 14 animals). L1 heat shock (N = 14 animals). L2 heat shock (N = 20 animals). L3 heat shock (N = 7 animals). L4 heat shock (N = 11 animals).

### Changes in AVA Membrane Potential Are Insufficient to Alter Presynaptic Organisation

K2P channels are well-established regulators of resting membrane potential (RMP)^13–16^, and our previous study demonstrated that TWK-40 activity significantly alters the steady-state membrane potential of AVA^23^. Thus, we hypothesised that the synaptic phenotypes of *twk-40* mutants might result from effects on membrane excitability.

To test this hypothesis, we first investigated the effect of mutations in other channels that alter the RMP. In particular, we examined mutations in sodium leak channels (*nca-1*/*nca-2*)^24,41,42^, whose loss hyperpolarises AVA^41^ and whose gain-of-function mutations enhance neuronal excitability^43^ (Figure 4A). Consistent with this role of Na^+^ leak in establishing membrane potential, we found that *nca-1(gf*) partially suppressed the locomotor phenotype of *twk-40(gf, L159N)* (Figure 4B), increased AVA neural activity as measured by calcium imaging (Figure 4C), and partially restored calcium transients observed in *twk-40(gf, L159N)* mutant animals (Figure 4C). However, despite this partial rescue of AVA activity and locomotor output, *nca-1(gf)* did not restore the altered SNB-1::GFP intensity (Figure 4D). These results suggest that the behavioural rescue and the presynaptic SNB-1 phenotype do not necessarily track together, and that TWK-40-dependent regulation of presynaptic organisation is unlikely to be explained solely by changes in AVA membrane potential or calcium activity.

**Figure 4.**
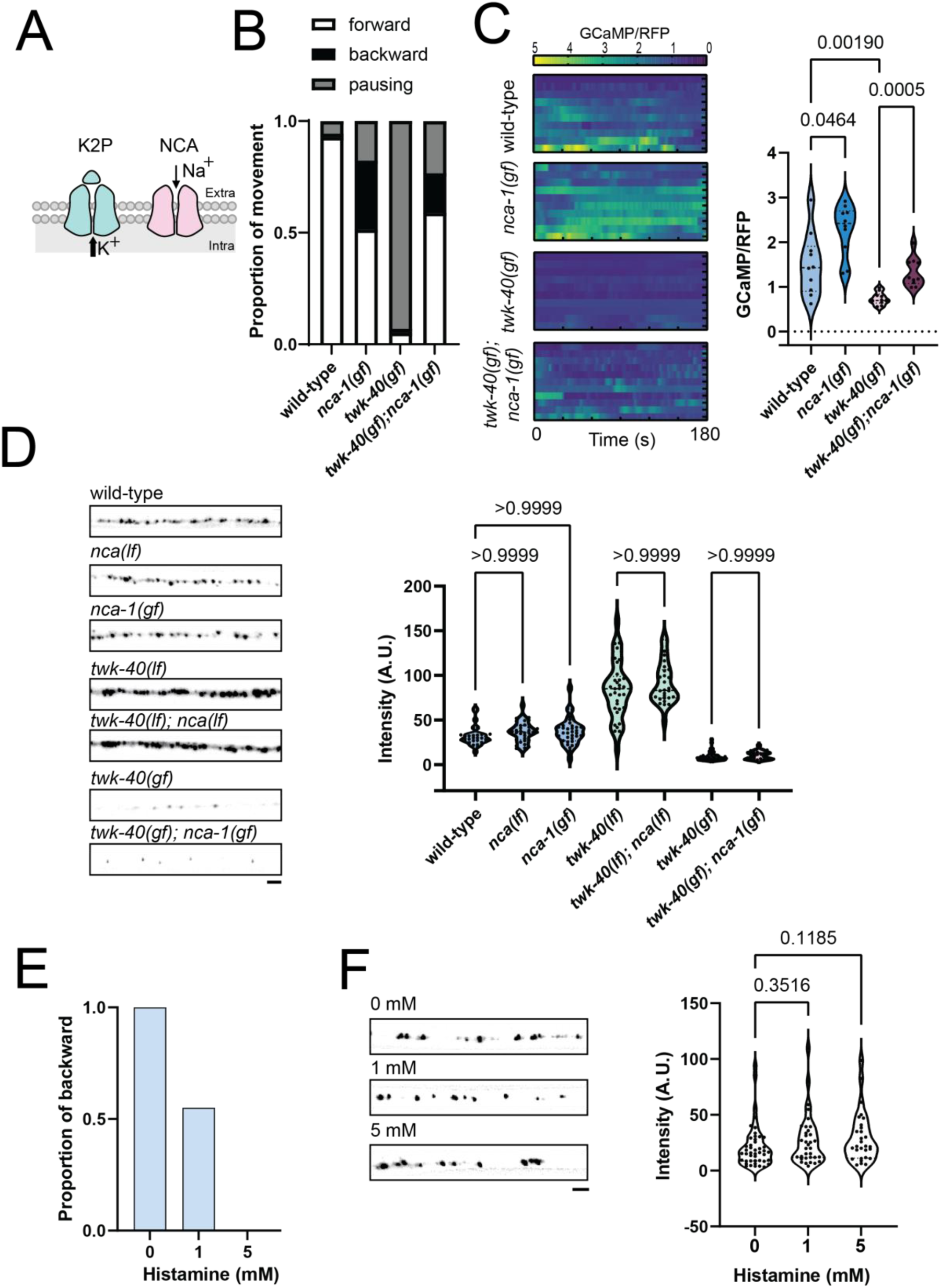
Membrane potential is not the sole regulator of presynaptic organisation. (A) Schematic illustration depicting how K2P (twk-40) and NCA (nca-1, nca-2) channels antagonise each other in regulating membrane potential. NCA-1 and NCA-2 function redundantly, requiring loss of both genes to disrupt NCA channel function, whereas gain-of-function of NCA-1 alone is sufficient to enhance channel activity43. (B) Proportion of movement (forward, backward, pausing) during spontaneous locomotion in each genotype, pooled across 10 animals per genotype. Gain-of-function in nca-1 partially suppresses the locomotor phenotype of twk-40(gf) animals. **Sample sizes:** N = 10 animals per genotype. (C) **Left:** Heatmap of GCaMP calcium traces in freely moving animals in each genotype, showing that nca-1(gf) partially rescues the reduced calcium activity observed in twk-40(gf) animals. **Right:** Quantification of average GCaMP/RFP ratio, showing significant differences across genotypes. **Sample sizes:** wild-type (N = 10 animals), nca-1(gf) (N = 10 animals), twk-40(gf) (N = 10 animals), twk-40(gf); nca-1(gf) (N = 10 animals). (D) **Left:** Representative confocal images of maximum intensity projections of AVA-SNB-1::GFP in each genotype. Scale bar: 2 μm. **Right:** Quantification of SNB-1::GFP intensity, showing that nca mutations do not significantly alter SNB-1::GFP levels in the corresponding genetic backgrounds, including twk-40 loss-and gain-of-function mutants. **Sample sizes:** wild-type (N = 31 animals), twk-40(lf) (N = 33 animals), twk-40(gf) (N = 51 animals), nca-2(lf); nca-1(lf) (nca(lf)) (N = 34 animals), nca-1(gf) (N = 34 animals), nca-2(lf) twk-40(lf); nca-1(lf) (N = 28 animals) and twk-40(gf); nca-1(gf) (N = 24 animals). (E) Backward probability upon head touch in animals expressing AVA-HisCl across different histamine concentrations. Backward movement is partially blocked at 1 mM histamine and fully blocked at 5 mM, indicating effective inhibition of AVA activity. Fisher’s exact test: 0 mM vs 1 mM histamine, P < 0.0001; 0 mM vs 5 mM histamine, P < 0.0001. **Sample sizes:** N = 80 animals per condition. (F) **Left:** Representative confocal images of maximum intensity projections of AVA-SNB-1::GFP under different histamine conditions. Scale bar: 2 μm. **Right:** Quantification of SNB-1::GFP intensity, showing no significant decrease across histamine treatments. **Sample sizes:** 0 mM (N = 45 animals), 1 mM (N = 34 animals), 5 mM (N = 29 animals).

To further test whether altered membrane excitability per se affected presynaptic organisation, we expressed a histamine-gated chloride channel in AVA to inhibit neuronal activity^44^. Histamine treatment blocked head touch-induced backward locomotion at 5 mM and partially blocked it at 1 mM after 1 h (Figure 4E). To assess the effects of prolonged treatment on presynaptic organisation, animals were maintained on histamine-containing plates for two generations. However, SNB-1::GFP intensity was not reduced at either concentration (Figure 4F), suggesting that reduced AVA excitability alone does not account for the presynaptic phenotype of twk-40(gf, L159N) mutants.

### The Presynaptic Effects of TWK-40 Scale with and Remain Genetically Coupled to Its Ion Channel Function

To determine whether the presynaptic effects of TWK-40 correlate with the severity of gain-of-function phenotypes, we compared two alleles carrying different substitutions at L159. The weaker gain-of-function allele, *twk-40(gf, L159T)*, is predicted to have lower channel activity than *twk-40(gf, L159N)*, based on studies of equivalent substitutions in homologous K2P channels^21^. Whereas the stronger gain-of-function allele, *twk-40(gf, L159N),* abolishes backward locomotion, mutants carrying the weaker allele, *twk-40(gf, L159T)*, exhibited a milder reduction in backward movement (Figure S4A). SNB-1::GFP levels in AVA were also reduced in *twk-40(gf, L159T)* mutants, but less severely than in *twk-40(gf, L159N)* animals (Figure S4B). Thus, across these two gain-of-function alleles, weaker TWK-40 channel activation was accompanied by milder locomotor and presynaptic phenotypes. Although this concordance suggested that the presynaptic phenotype tracks TWK-40 channel function, it did not establish whether the two effects could be genetically separated.

We next searched for a separation-of-function construct that might dissociate the presynaptic phenotype from the canonical effects of TWK-40 channel activity. To this end, we generated truncated TWK-40(gf, L159N) proteins and expressed them specifically in AVA (Figure S4C). Across the constructs tested, truncations that retained the gain-of-function locomotor phenotype also retained the presynaptic defect, whereas truncations that lost the locomotor phenotype also lost the presynaptic defect (Figure S4D, E). We therefore did not identify a construct that genetically separated these two phenotypes. Together, the allelic and truncation analyses indicate that the presynaptic effects of TWK-40 track, and within the genetic perturbations tested remain closely coupled to, its canonical ion channel function. However, these experiments do not establish whether the presynaptic effects of TWK-40 are directly caused by its ion channel activity.

### Intracellular Potassium Homeostasis Is Essential for Presynaptic Organisation

K2P channels influence both membrane potential and intracellular K⁺ homeostasis^45–48^. As the presynaptic phenotype could not be explained solely by altered AVA excitability or membrane potential, we hypothesised that TWK-40 might regulate presynaptic organisation through effects on intracellular potassium homeostasis. To test this, we first examined whether intracellular K^+^ levels were indeed altered in *twk-40* mutant animals using Kirin (Figure 5A), a genetically encoded Förster resonance energy transfer (FRET)-based potassium sensor^46,49^. We observed an increased Kirin YFP/CFP ratio in *twk-40(lf)* mutants and a decreased ratio in *twk-40(gf, L159N)* mutants (Figure 5A), consistent with increased and decreased intracellular K⁺ levels, respectively. These results correspond to the reduced and increased K^+^ efflux expected in the loss- and gain-of-function mutants, respectively.

**Figure 5.**
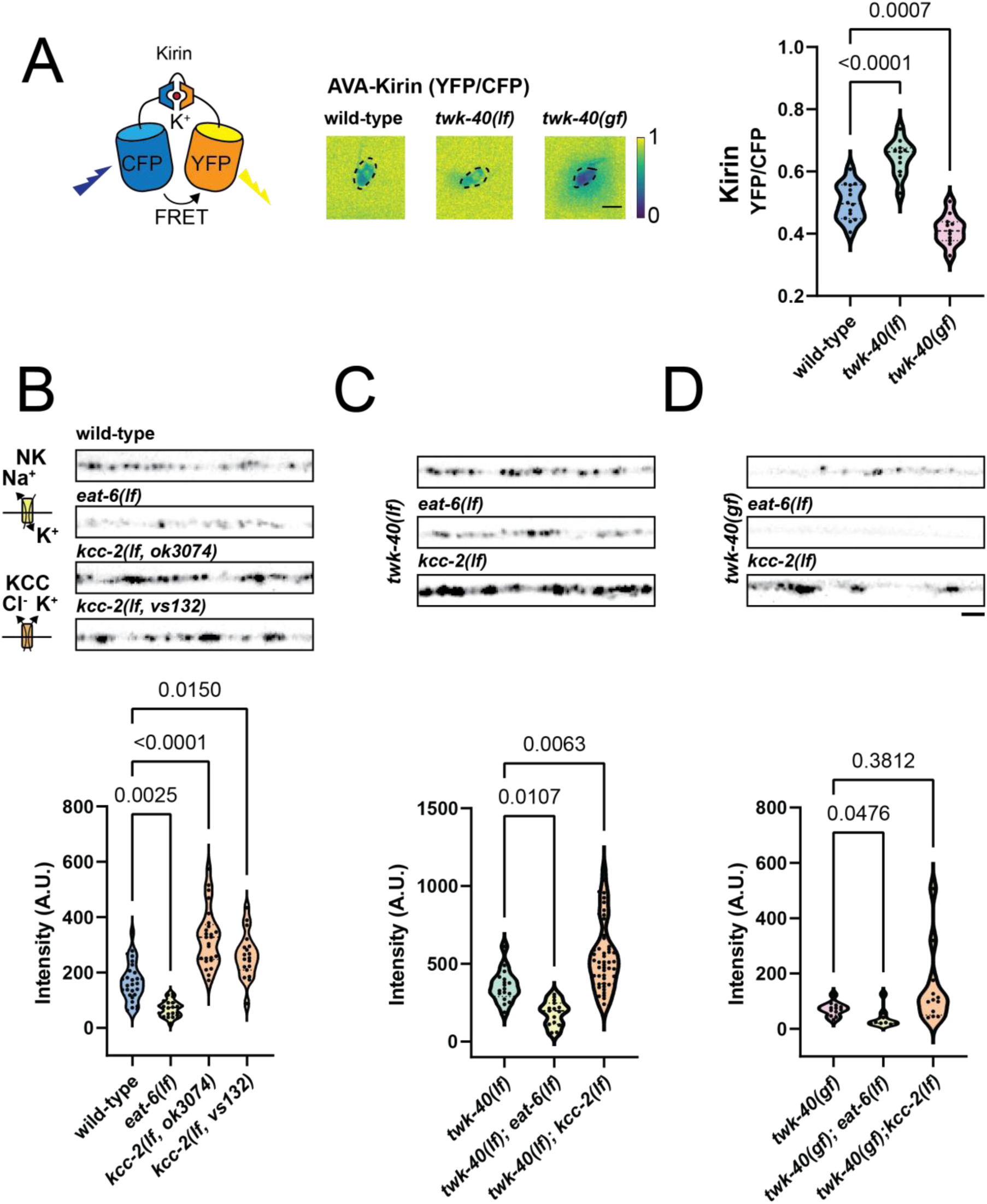
Intracellular potassium homeostasis regulated by TWK-40 is essential for proper presynaptic organisation. (A) **Left:** Schematic illustration of Kirin, a FRET-based potassium sensor used to measure intracellular K⁺ levels. Middle: Representative images of YFP/CFP fluorescence ratio in AVA neurons. **Right:** Quantification of YFP/CFP fluorescence ratio in AVA neurons, showing a significant increase in twk-40(lf) and a significant decrease in twk-40(gf). **Sample sizes:** wild-type (N = 14 animals), twk-40(lf) (N = 11 animals), twk-40(gf) (N = 11 animals). Scale bar: 5 μm. (B) **Top:** Representative confocal images showing SNB-1::GFP intensity in AVA of wild-type, eat-6(lf), and kcc-2(lf). **Bottom:** Quantification of SNB-1::GFP intensity, showing a significant decrease in eat-6(lf) and an increase in kcc-2(lf). **Sample sizes:** wild-type (N = 28 animals), eat-6(lf) (N = 22 animals), kcc-2(lf, ok3074) (N = 27 animals), kcc-2(lf, vs132) (N = 19 animals). (C) **Top:** Representative confocal images showing SNB-1::GFP intensity in AVA in the twk-40(lf) background, with eat-6(lf) showing decreased SNB-1 intensity and kcc-2(lf) showing increased intensity. **Bottom:** Quantification of SNB-1::GFP intensity, showing a significant reduction in twk-40(lf); eat-6(lf) and an increase in twk-40(lf); kcc-2(lf). **Sample sizes:** twk-40(lf) (N = 17 animals), twk-40(lf); eat-6(lf) (N = 35 animals), twk-40(lf); kcc-2(lf) (N = 51 animals). (D) **Top:** Representative confocal images showing SNB-1::GFP intensity in AVA in twk-40(gf) background, with eat-6(lf) showing decreased intensity and kcc-2(lf) showing no significant increase. **Bottom:** Quantification of SNB-1::GFP intensity, showing a modest but significant decrease following eat-6(lf), whereas kcc-2(lf) showed a non-significant trend towards increased intensity in the twk-40(gf) background. **Sample sizes:** twk-40(gf) (N = 12 animals), twk-40(gf); eat-6(lf) (N = 15 animals), twk-40(gf); kcc-2(lf) (N = 12 animals). (B–D) Scale bar: 2 μm.

To assess whether these changes in intracellular potassium levels could regulate presynaptic organisation, we tested whether other genes affecting intracellular potassium concentrations, in particular potassium transporters, could exert similar effects on synaptic vesicle proteins. We first analysed mutants defective in *kcc-2,* which encodes a potassium-chloride cotransporter^50^, and whose loss, like loss of K2P channel activity, should elevate intracellular K^+^ concentration. Indeed, we observed that SNB-1::GFP expression was significantly increased in *kcc-2(lf)* mutants, similar to the phenotype of *twk-40(lf)* (Figure 5B). We also analysed mutations in *eat-6,* which encodes a sodium-potassium pump α-subunit ^51^. Loss-of-function mutations in *eat-6* would be expected to reduce the intracellular K^+^ concentration, similar to a K2P gain-of-function mutation, but depolarise rather than hyperpolarise the membrane. In fact, we observed a significant reduction in SNB-1::GFP intensity in *eat-6(lf)* (Figure 5B), consistent with a role for intracellular potassium homeostasis in the presynaptic phenotype of *twk-40(gf, L159N)* animals.

Transporter and K2P channel double mutants further supported the hypothesis that intracellular K⁺ levels influence TWK-40-dependent presynaptic organisation. In the *twk-40(lf)* background, *kcc-2(lf)* further increased SNB-1::GFP levels, whereas *eat-6(lf)* significantly reduced them (Figure 5C). In the *twk-40(gf, L159N)* background, *eat-6(lf)* further reduced SNB-1::GFP intensity, although the effect was modest, whereas *kcc-2(lf)* showed a trend towards increased intensity that did not reach statistical significance (Figure 5D). Together, these results further support a role for potassium homeostasis in regulating synaptic organisation.

### The Synaptic Effects of TWK-40 Depend on a Potassium-Sensitive Enzyme

We reasoned that the effects of intracellular potassium concentrations on synapses might depend on potassium-sensitive effector proteins. One candidate we considered was pyruvate kinase (PYK-1), a glycolytic enzyme with conserved residues implicated in K⁺-dependent catalysis^52^. Mammalian pyruvate kinase has also been shown to “moonlight” as a protein kinase^53^. When we overexpressed *pyk-1* using a genomic transgene containing endogenous regulatory sequences in a wild-type background, we observed increased SNB-1::GFP intensity in AVA neurons (Figure 6A). Overexpressing a mutant PYK-1 variant (T121L) (Figure 6B), analogous to a rabbit mutation (T120L)^52^, which targets a residue associated with K⁺-dependent catalytic activity^52^, failed to enhance SNB-1::GFP levels in wild-type animals (Figure 6A). Thus, the conserved K⁺-associated residue appears to be important for the ability of PYK-1 to promote presynaptic SNB-1::GFP accumulation.

**Figure 6.**
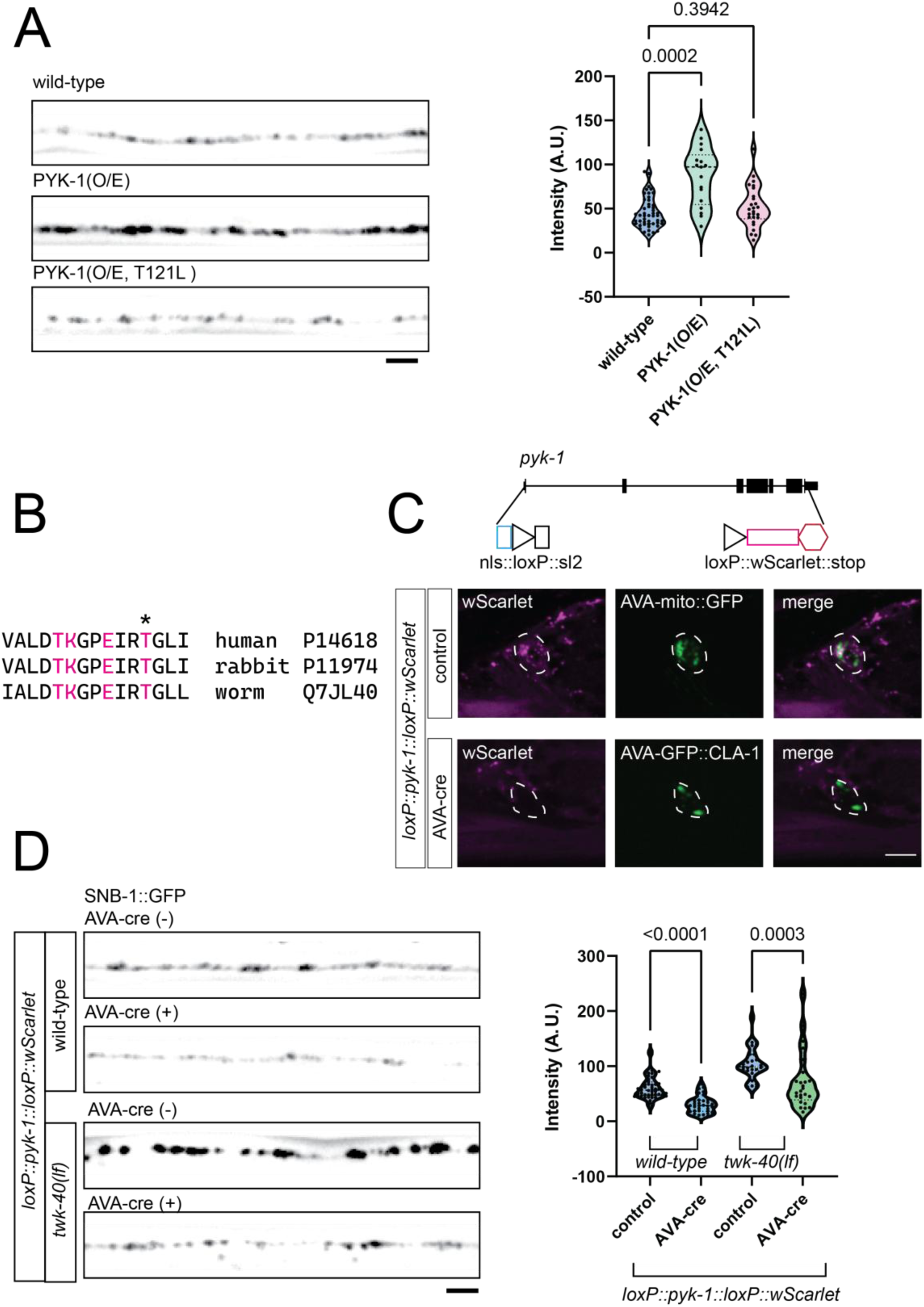
PYK-1 Regulates Presynaptic Organisation. (A) **Left:** Representative confocal images of SNB-1::GFP intensity in AVA neurons for each genotype. Scale bar: 2 μm. **Right:** Quantification of SNB-1::GFP intensity, showing a significant increase in PYK-1 overexpression (O/E) but not in PYK-1(T121L) (O/E) animals. **Sample sizes:** wild-type (N = 48 animals), PYK-1(O/E) (N = 17 animals), PYK-1T121L(O/E) (N = 26 animals). The mutated residue corresponds to a conserved site associated with K⁺-dependent pyruvate kinase activity. (B) Sequence alignment of the region surrounding the conserved K⁺-dependent pyruvate kinase signature in human PKM (UniProt: P14618), rabbit PKM (UniProt: P11974), and C. elegans PYK-1 (UniProt: Q7JL40). Residues associated with K⁺-dependent pyruvate kinase activity are highlighted in magenta. The asterisk marks the threonine mutated to leucine at position 121 in this study. Rabbit residue numbering follows the convention used in the cited biochemical literature, which excludes the initiator methionine. (C) **Top:** Schematic of the CRISPR/Cas9-engineered pyk-1 locus with the pyk-1a coding region flanked by loxP sites, allowing Cre-mediated conditional deletion of pyk-1. **Bottom:** Representative confocal images showing expression of pyk-1::wScarlet in control animals (top row) and loss of signal in AVA-cre animals (bottom row), consistent with Cre-mediated deletion of pyk-1 in AVA neurons. AVA neurons are labelled with AVA-mito::GFP or AVA-GFP::CLA-1. Cre-mediated recombination excises the loxP-flanked pyk-1 genomic region, resulting in the loss of PYK-1::wScarlet fluorescence in AVA. Dashed lines denote AVA. Scale bar: 5 μm. (D) **Left:** Representative confocal images of SNB-1::GFP intensity in AVA neurons for each genotype. Scale bar: 2 μm. **Right:** Quantification of SNB-1::GFP intensity, showing a significant decrease in wild-type and twk-40(lf) background animals. **Sample sizes:** wild-type control (N = 51 animals), wild-type, AVA-cre (N = 36 animals), twk-40(lf) control (N = 27 animals), twk-40(lf), AVA-cre (N = 26 animals).

To investigate the contribution of PYK-1 to presynaptic organisation in AVA, we engineered a conditional knockout allele by inserting loxP sites flanking the endogenous *pyk-1* locus, with wScarlet fused to its C-terminus (Figure 6C top). Cre expression resulted in the loss of PYK-1::wScarlet fluorescence in AVA, consistent with successful Cre/loxP-mediated conditional deletion (Figure 6C). We observed a significant reduction in AVA SNB-1::GFP intensity following conditional deletion (Figure 6D). Furthermore, this manipulation reduced the elevated SNB-1::GFP levels in twk-40(lf) animals (Figure 6D). These results support a role for PYK-1 in presynaptic organisation in AVA and suggest that intracellular potassium levels could regulate the synapse through a PYK-1-dependent pathway.

### Transcriptional Pathways Involved in Synaptic Maintenance

To identify additional downstream players involved in potassium-dependent synapse development, we carried out transcriptomic comparisons between wild-type and *twk-40* mutant animals (Figure S7A, Methods). We performed bulk RNA sequencing (RNA-seq) on fluorescence-activated cell sorting (FACS)-isolated TWK-40-expressing neurons, AVA included, and compared the transcriptional profiles of wild-type, *twk-40(lf)*, and *twk-40(gf, L159N)* mutant animals. Between wild-type and *twk-40(lf)* mutants, 44 genes met our thresholds for differential expression (Figure S7B; Table S2). In contrast, 776 genes met these thresholds in *twk-40(gf, L159N)* mutants, with most differentially expressed genes (DEGs) downregulated and distinct expression patterns across biological replicates (Figure 7A, B and Figure S7D; Table S1). GO enrichment analysis using the RNA-seq-derived reference set identified significant enrichment of biological processes associated with synaptic organisation and transmission, potassium ion transport, axon guidance, and neuronal differentiation (Figure 7C and Figure S7C; Tables S3–S5). The differentially expressed genes also included genes encoding ion channels (including K2Ps), neuropeptides, transcription factors, and extracellular matrix proteins (Table S1). Among these downregulated genes were *cla-1*, encoding the active zone protein CLA-1 examined in our initial presynaptic marker analysis (Figure 2), as well as *snt-4*, encoding a synaptotagmin homolog. SNT-4::GFP fluorescence was also reduced in *twk-40(gf, L159N)* animals (Figure S7E). Thus, increased TWK-40 activity is associated with reduced expression of genes involved in synaptic organisation and function.

**Figure 7.**
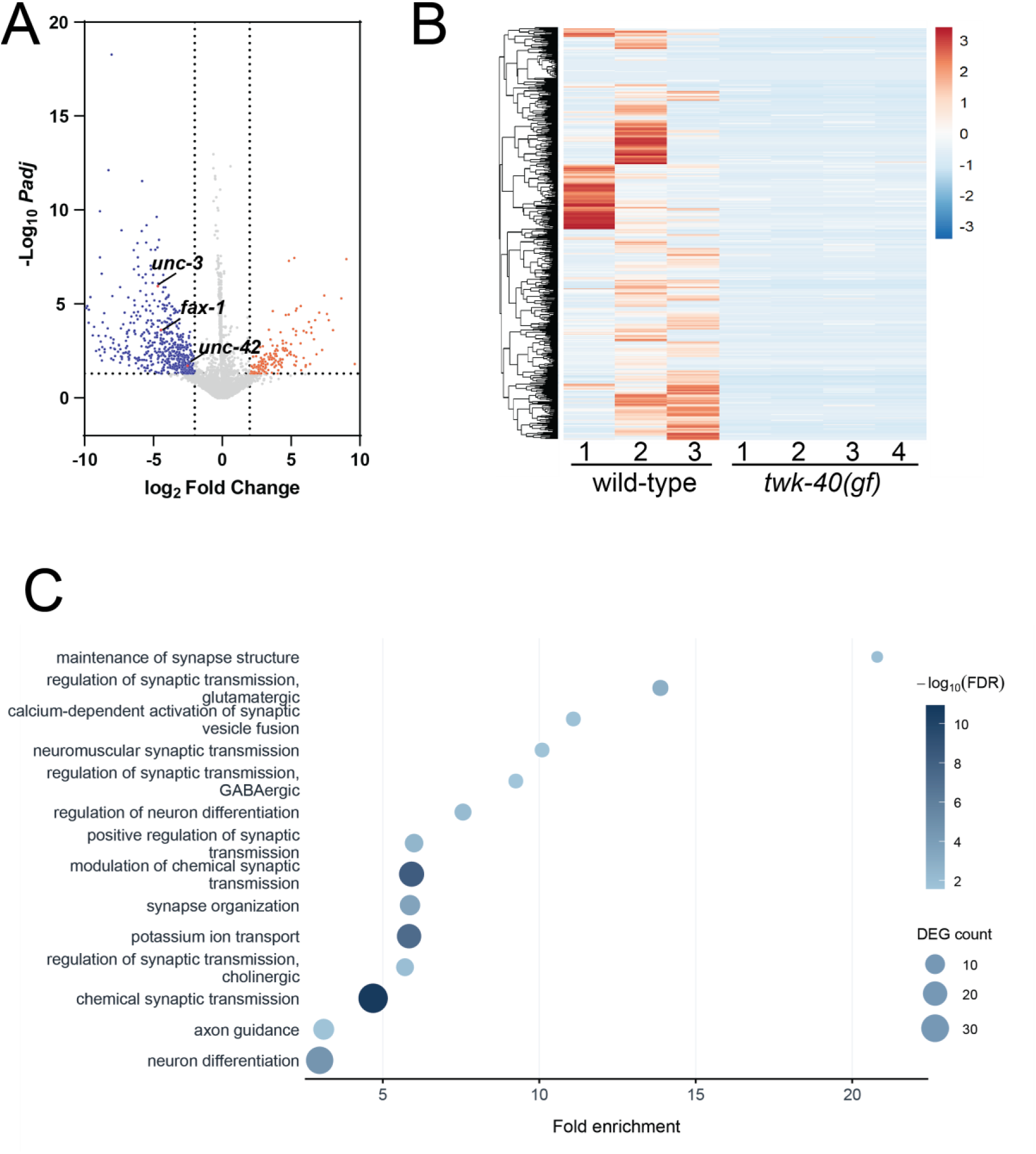
twk-40(gf) alters neuronal transcriptional programs, including genes related to presynaptic function. (A) Volcano plot showing differentially expressed genes in twk-40-expressing neurons from twk-40(gf) animals. Notable genes (unc-3, fax-1, unc-42) with significant downregulation are highlighted; the corresponding DEG list is provided in Table S1. **Sample size:** wild-type control (N = 3 biological replicates); twk-40(gf) (N = 4 biological replicates). (B) Heatmap showing row-scaled, variance-stabilised expression values for the top 500 differentially expressed genes in twk-40(gf) animals. Genes were hierarchically clustered, whereas samples were arranged by genotype without column clustering. Each column represents an individual biological replicate (wild-type, N = 3; twk-40(gf), N = 4). (C) Gene Ontology (GO) biological process overrepresentation analysis of differentially expressed genes in twk-40(gf) animals using an RNA-seq-derived reference set. Fourteen significantly enriched GO terms related to synaptic organisation, neuronal function, and potassium ion transport are shown. The x-axis represents fold enrichment, dot size indicates the number of differentially expressed genes associated with each GO term, and colour represents −log₁₀(FDR) (Tables S3 and S5).

We also identified a small set of transcriptional regulators that were significantly downregulated in *twk-40(gf, L159N)* mutants (Figure 7A). These included *unc-3* (log₂FC = −4.69, adjusted P = 1.11 × 10⁻⁶), encoding UNC-3, a homolog of mammalian Olf-1/Early B-cell Factor proteins and a terminal selector for cholinergic motor neuron differentiation^54,55^; *unc-42* (log₂FC = -2.52, adjusted P = 0.0188), encoding UNC-42, a homeodomain transcription factor essential for interneuron identity^56–58^; and *fax-1* (log₂FC = -4.46, adjusted P = 0.0002), encoding FAX-1, a nuclear hormone receptor associated with axonal differentiation^58^. Protein-fusion reporters confirmed reduced UNC-3 and FAX-1 levels in *twk-40(gf, L159N)* mutants, whereas UNC-42::GFP fluorescence showed a downward trend that did not reach statistical significance (Figure 8A, B). Conversely, UNC-3::EGFP and UNC-42::GFP reporters were significantly increased in *twk-40(lf)* mutants (Figure 8A, B). The *unc-42* transcript was also upregulated in *twk-40(lf)* (log₂FC = +2.08, adjusted P = 0.0411; Table S2).

**Figure 8.**
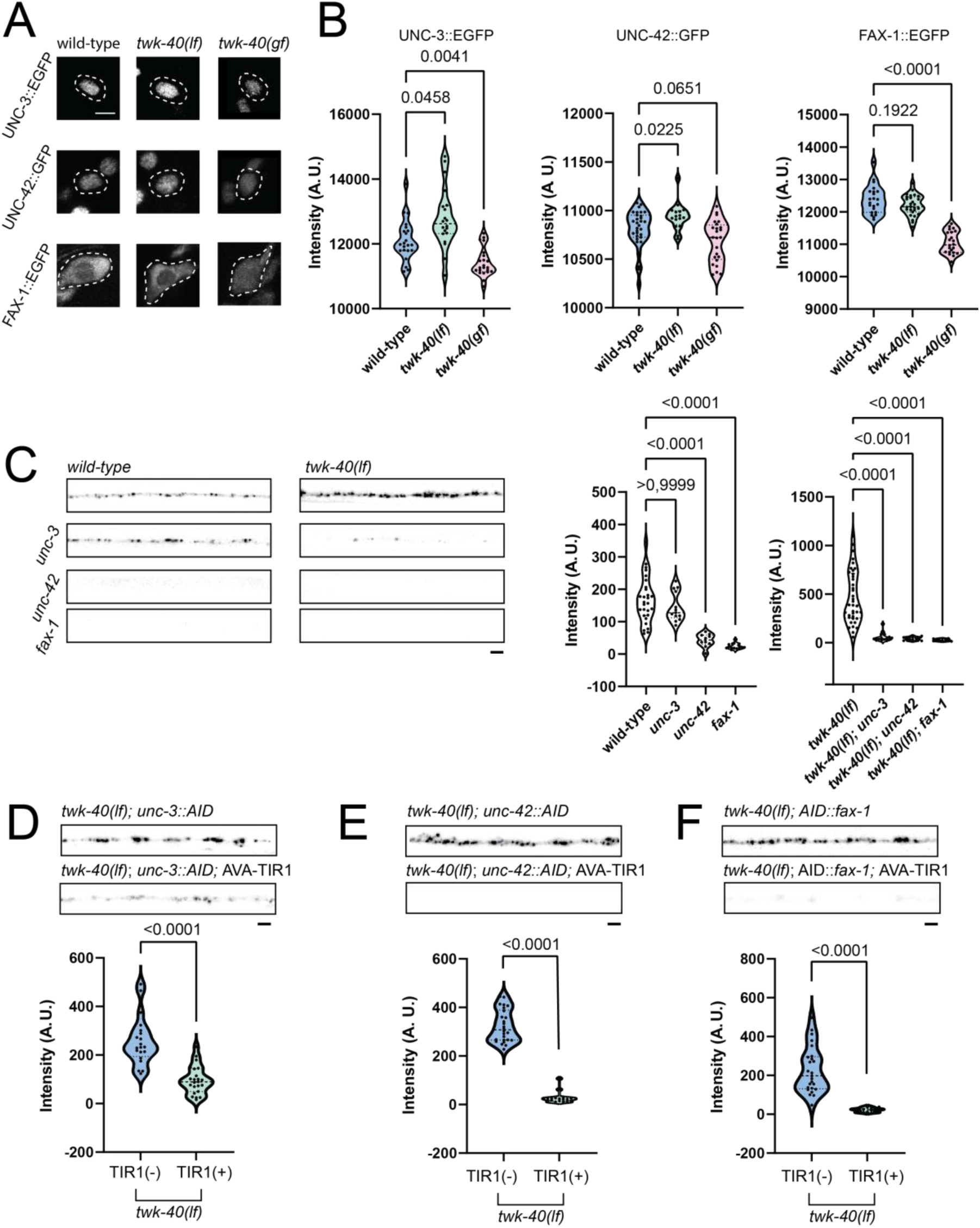
UNC-3, UNC-42, and FAX-1 contribute to TWK-40-dependent regulation of presynaptic SNB-1::GFP levels. (A) Representative confocal images of UNC-3::EGFP, UNC-42::GFP, and FAX-1::EGFP fluorescence in AVA neurons across different genotypes (wild-type, twk-40(lf), and twk-40(gf)). UNC-3::EGFP and FAX-1::EGFP levels are decreased in twk-40(gf), whereas UNC-42::GFP shows a non-significant downward trend. Dashed outlines denote AVA. Scale bar: 1 μm. (B) Quantification of UNC-3::EGFP, UNC-42::GFP, and FAX-1::EGFP fluorescence intensity in AVA neurons. **Sample sizes (unc-3 reporter group):** wild-type (N = 22 animals), twk-40(lf) (N = 22 animals), twk-40(gf) (N = 20 animals). **Sample sizes (unc-42 reporter group):** wild-type (N = 31 animals), twk-40(lf) (N = 16 animals), twk-40(gf) (N = 24 animals). **Sample sizes (fax-1 reporter group):** wild-type (N = 22 animals), twk-40(lf) (N = 27 animals), twk-40(gf) (N = 22 animals). (C) **Left:** Representative confocal images of AVA-SNB-1::GFP fluorescence along the ventral nerve cord in wild-type, unc-3(lf), unc-42(lf), fax-1(lf), and corresponding twk-40(lf) double mutants. Loss of unc-42 or fax-1 reduces presynaptic SNB-1::GFP levels in wild-type. Loss of unc-3, unc-42, or fax-1 reduces presynaptic SNB-1::GFP levels in twk-40(lf). **Right:** Quantification of SNB-1::GFP fluorescence intensity. **Sample sizes:** wild-type (N = 28 animals), unc-3(lf) (N = 13 animals), unc-42(lf) (N = 10 animals), fax-1(lf) (N = 14 animals), twk-40(lf) (N = 38 animals), twk-40(lf); unc-3(lf) (N = 16 animals), twk-40(lf); unc-42(lf) (N = 13 animals), twk-40(lf); fax-1(lf) (N = 10 animals). (D–F) AVA-specific auxin-inducible degradation of UNC-3 (D), UNC-42 (E), or FAX-1 (F) in twk-40(lf) animals. **Top:** Representative confocal images of AVA-SNB-1::GFP fluorescence. **Bottom:** Quantification of SNB-1::GFP fluorescence intensity, showing a significant reduction in animals expressing AVA-specific TIR1 compared with the corresponding TIR1-negative controls. In each experiment, both groups carried the corresponding AID-tagged transcription factor in a twk-40(lf) background, and all animals were treated with 4 mM K-NAA. TIR1 (TRANSPORT INHIBITOR RESPONSE 1) is an auxin receptor that mediates degradation of AID-tagged proteins. **Sample sizes:** (D) UNC-3::AID: TIR1(−) (N = 20 animals), TIR1(+) (N = 23 animals). (E) UNC-42::AID: TIR1(−) (N = 26 animals), TIR1(+) (N = 19 animals). (F) AID::FAX-1: TIR1(−) (N = 25 animals), TIR1(+) (N = 20 animals). Scale bars: 2 μm.

To assess the potential role of these transcription factors in presynaptic organisation, we assayed SNB-1::GFP expression in *unc-3*, *unc-42*, and *fax-1* loss-of-function mutants. We observed reduced AVA SNB-1::GFP reporter fluorescence in *unc-42* and *fax-1* mutants, including in the *twk-40(lf)* background (Figure 8C). To assess their contribution within AVA, we used the auxin-inducible degradation (AID) system^59,60^ to target UNC-3, UNC-42, or FAX-1 in a *twk-40(lf)* background. SNB-1::GFP intensity was reduced in all three experimental groups compared with their corresponding TIR1-negative controls (Figure 8D–F). Together, these data identify UNC-3, UNC-42, and FAX-1 as transcriptional regulators that are downregulated in *twk-40*(gf, L159N) neurons and contribute to presynaptic SNB-1::GFP levels.

## Discussion

We have shown here that TWK-40, a K2P channel previously shown to play an essential role in setting the membrane potential of *C. elegans* interneurons, also contributes to the maintenance of presynaptic organisation. Our findings suggest that the presynaptic effects of TWK-40 are linked to intracellular potassium homeostasis and cannot be explained solely by changes in membrane potential. We also identify multiple downstream regulatory layers, including the potassium-sensitive enzyme PYK-1 and transcriptional regulators identified through candidate screening. Together, these findings strongly support a role for intracellular potassium homeostasis in TWK-40-dependent presynaptic organisation, beyond the effects of TWK-40 on membrane potential (Figure 9).

**Figure 9.**
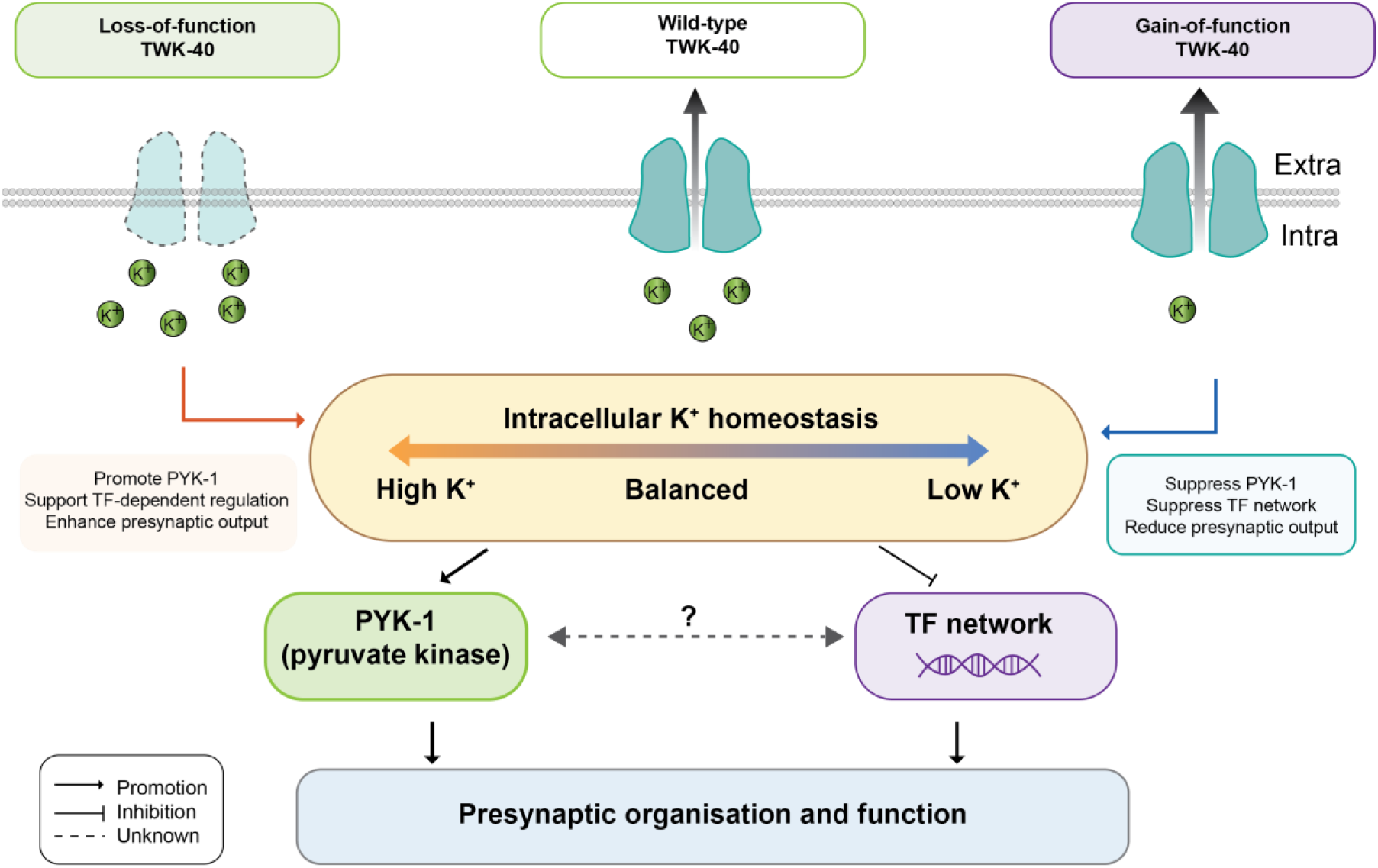
A model for TWK-40-dependent regulation of presynaptic organisation through intracellular K⁺ homeostasis. Altered TWK-40 activity shifts intracellular potassium balance, which modulates PYK-1 and a transcriptional network to control presynaptic organisation and function.

### Potassium Homeostasis as a Regulator of Synaptic Structure

A major insight from our study is that TWK-40’s role in presynaptic organisation is independent of its canonical function in regulating membrane potential, but correlates instead with its effect on potassium homeostasis. While mutations in *twk-40* strongly affect the resting membrane potential of the AVA neurons, other mutations with similar effects on the membrane potential, such as in sodium leak channels, do not replicate TWK-40’s synaptic phenotypes. In contrast, mutations in other genes that elevate or reduce intracellular potassium levels (e.g. K^+^ pumps) show similar synaptic phenotypes to mutations in *twk-40* with similar effects on K^+^ concentrations. Most notably, loss-of-function mutations in the sodium/potassium ATPase *eat-6,* expected to lower intracellular K^+^ and depolarise the membrane, reduced the abundance of synaptic proteins, similar to *twk-40(gf)* mutations that lower intracellular K^+^ but hyperpolarise the membrane. Together, these results support a major contribution of intracellular potassium homeostasis to the synaptic phenotypes of *twk-40* mutants, beyond effects attributable to membrane potential.

This unexpected connection between potassium homeostasis and presynaptic organisation highlights a potentially unappreciated mechanism by which ion channels could contribute to neural circuit stability, distinct from effects on membrane potential. While the efflux through potassium leak channels is usually thought to be minimal when the RMP is close to potassium equilibrium potential (E_K_), in neurons with a more depolarised resting membrane (such as AVA), the efflux through a K2P channel could be significant enough to affect intracellular concentrations. Indeed, potassium channels such as TASK2/KCNK5, TWIK2/KCNK6, and TREK-1/KCNK2 have been implicated in K⁺ efflux or K⁺-dependent signalling in diverse cell types and organisms^45–48^. Interestingly, various reports from *C. elegans* and other systems are consistent with a link between potassium homeostasis and neuronal differentiation, cytoskeletal dynamics, and synaptic maturation, though these effects have often been interpreted in the context of membrane depolarisation or altered excitability, rather than intracellular K⁺ homeostasis per se^61–67^. For example, mutations in the K⁺/Cl⁻ cotransporter KCC2 have also been linked to synaptic structure in both invertebrate and vertebrate systems^50,68^. Together, these data support a model in which potassium homeostasis is a direct signal that can regulate synapse assembly and maintenance.

Interestingly, we observed that the *C. elegans* K2P channel TWK-40 is enriched at SNB-1-positive presynaptic regions along the ventral nerve cord. K2P channels in vertebrate neurons can exhibit subcellular compartmentalisation as well. For example, TRAAK and TREK-1 localise to nodes of Ranvier, where they support conduction by regulating local K⁺ dynamics^18^. Such spatial restriction of K⁺ flux raises the possibility that localised potassium gradients may influence structural aspects of neurons, such as presynaptic assembly or cytoskeletal architecture.

### Pyruvate Kinase as a Potential Mediator of Potassium-Dependent Synaptic Regulation

The molecular mechanisms that could link intracellular potassium to the regulation of synaptic function remain largely unexplored. In this study, we identified PYK-1, a potassium-dependent enzyme, as a candidate downstream effector in the TWK-40 pathway regulating presynaptic organisation. Loss of PYK-1 mimics the reduced-synaptic SNB-1 phenotype of mutants with low intracellular potassium and suppresses the increased-synaptic phenotype caused by loss of TWK-40. Conversely, overexpression of PYK-1 increases AVA SNB-1::GFP levels, whereas the T121L variant, which alters a conserved residue important for pyruvate kinase catalytic activity, does not. Together, these results support a role for PYK-1 in presynaptic organisation and suggest that its enzymatic function may contribute to the effects of TWK-40-dependent potassium homeostasis. In the future, we hope to identify such downstream effectors and elucidate the causal relationship between potassium dynamics and PYK-1 activity in neurons.

How could PYK-1 influence synaptic function? Pyruvate kinase plays a critical role in glycolysis, catalysing the final step through which ATP is generated by the transfer of phosphate from phosphoenolpyruvate to ADP. Vesicles bearing synaptic proteins and glycolytic enzymes, potentially corresponding to synaptic vesicle precursors, have been shown to generate ATP locally to sustain axonal transport^69–71^. This raises the possibility that PYK-1 may contribute to synaptic organisation by supporting local ATP generation or vesicle trafficking. Notably, hippocampus-specific depletion of PKM2 causes cognitive defects that can be rescued by lactate, a product of pyruvate metabolism, consistent with a metabolic link between pyruvate kinase activity and neural development^72^.

Alternatively, pyruvate kinase, like several other glycolytic enzymes, has been shown to “moonlight” as a protein kinase in a variety of cell types^53^. For example, in mammals the PKM2 isoform of pyruvate kinase can translocate to the nucleus under specific conditions, where it modulates gene expression by phosphorylating histone H3^73^, thus acting as a transcriptional co-regulator^74^. Nuclear PKM2 has also been reported to phosphorylate STAT3^75^ and to interact with Oct4^76^ thereby influencing transcriptional regulation. PKM2 has also been shown to phosphorylate protein substrates in the cytoplasm, for example the TOR pathway component AKT1S1^77^. Thus, PYK-1 could in principle play a direct role in mediating the transcriptional changes observed in *twk-40(gf)* neurons, or alternatively play a more indirect role through its effects on glycolytic metabolism. Determining how PYK-1-dependent metabolic regulation intersects with transcriptional control of synaptic organisation in *C. elegans* could serve as a tractable model for similar and potentially conserved processes in other organisms.

### Terminal Selector-Like Regulation of Synaptic Organisation

Our findings suggest that transcriptional regulation of presynaptic organisation in AVA may differ from previously described activity-dependent programmes. In *C. elegans* dopaminergic neurons, for example, recent work has shown that synaptogenesis is promoted through an activity-regulated transcriptional program involving EGL-43/MECOM and FOS-1/FOS, which directly activate presynaptic genes in coordination with terminal selectors and CUT homeobox transcription factors^4^. This pathway parallels activity-dependent mechanisms of synaptic gene regulation described in other organisms, including vertebrates^5,78,79^. However, such activity-dependent mechanisms may not be universal in all neuron classes. Many neurons form presynaptic structures before the onset of action potentials or synaptic activity^80,81^. Indeed, the cholinergic premotor interneuron AVA, the focus of this study, exhibits tonic, action potential-independent activity and functions within the core locomotor command circuit of *C. elegans*. Thus, these neurons might be expected to adopt different gene regulation strategies than dopaminergic neurons.

Our findings suggest that terminal-selector-like transcriptional regulation contributes to presynaptic organisation in AVA neurons. Terminal selectors are transcription factors that define and maintain neuronal identity by directly activating gene expression programs for neurotransmitter synthesis, ion channel function, and synaptic organisation. In *C. elegans*, terminal selectors such as UNC-86^82^, TTX-3^82^, and UNC-42^57^ have been shown to coordinate neuron-type-specific expression of effector genes throughout the lifespan. Our findings suggest that potassium, regulated by TWK-40, is associated with the expression of transcription factors that function similarly to terminal selectors in stabilising presynaptic organisation. Activity-regulated transcription factors such as CREB^83^, MEF2^6^, and FOS^7^ also contribute to synaptic gene expression and plasticity^84^. Thus, cholinergic premotor interneurons may use terminal selector-like transcriptional programs to maintain presynaptic organisation^55,85^. These programmes may provide a stable, intrinsic framework for synaptic organisation, consistent with developmental connectomic studies showing that core interneuron circuits remain relatively stable while sensory and motor pathways undergo substantial remodelling^22^. This suggests that different neuronal classes may favour distinct strategies for synaptic maintenance: sensory neurons prioritising adaptability and activity-dependent plasticity, and premotor interneurons emphasising stability and precision^4,22^.

## Author Contributions

J.M. conceived the study, developed the overall research strategy and experimental design, performed the majority of the experiments, including synaptic imaging, behavioural assays, optogenetic experiments, cell sorting, and calcium imaging, analysed and interpreted the data, and wrote the original draft of the manuscript. N.R. performed experiments for RNA sequencing and carried out the RNA-seq data analysis. N.R. and Y.L. performed a subset of the synaptic imaging experiments and cell sorting. J.M., N.R., Y.L., and T.B. generated and provided strains and reagents used in the study. W.S., M.Z., and I.B. contributed to data interpretation and critical revision of the manuscript. W.S. supervised the study, contributed to data interpretation, and contributed to writing the manuscript. All authors reviewed and edited the manuscript.

## Acknowledgements

We thank Ying Wang and Wesley Hung (Lunenfeld-Tanenbaum Research Institute) for providing reagents and technical support; Kin Chan (Lunenfeld-Tanenbaum Research Institute) for assistance with RNA sequencing and the University of Toronto Flow Cytometry Facility for assistance with cell sorting; John Jin (Lunenfeld-Tanenbaum Research Institute) for IT support; the Light Microscopy Facility (MRC Laboratory of Molecular Biology) and the VIB Bioimaging Core Leuven (VIB, Leuven) for technical support and access to imaging instrumentation; and Christopher L. Antos and Yingchuan Qi (ShanghaiTech University) for providing reagents; and Denise Walker, Isabel Conze, and Joe Gehler-Rahman (MRC Laboratory of Molecular Biology) for proofreading. J.M. also thanks Sadaharu Meng for his companionship throughout this project.

## Funding

This work was supported by the Research Foundation Flanders (FWO; V440424N to J.M. and G079521N to W.S.), the Irving R. Gerstein Research Fellowship (to J.M.), the KU Leuven Research Council (C16/25/005 to I.B.), the Baillet Latour Fund (to I.B.), the China Scholarship Council (CSC) Scholarship (to Y.L.), the National Natural Science Foundation of China (32371189 to S.G. https://www.nsfc.gov.cn/), the European Research Council (*Kelegans* to T.B.), the Medical Research Council (UK; MC-M102-M1421 to W.S.), the Canadian Institutes of Health Research, the Natural Sciences and Engineering Research Council of Canada (NSERC; RGPIN-2024-06020), and the Lundbeck Foundation (all to M.Z.).

## Materials and Methods

### Constructs and strains

A list of constructs and strains used in this study is provided in Table S6. Constructs and strains are available upon request. All *C. elegans* strains were cultured on the standard Nematode Growth Medium (NGM) plates seeded with *Escherichia coli* OP50 and maintained at 22°C unless specified. The wild-type animal refers to the Bristol N2 strain or control animals as specified in the figure legends. L4-stage hermaphrodites were used for all experiments unless specified otherwise.

### Isolation, cloning, and generation of K2P mutants

Strains *hp834* [*twk-40(lf)*], *bln336* [*twk-40(gf, L159N)*], and *bln282* [*twk-40*::tagRFP::ZF] were previously engineered using CRISPR/Cas9^21,23^. Unless otherwise indicated, *twk-40(gf)* refers to the L159N allele.

The weak gain-of-function allele *twk-40*(*gf*, *L159T)* was generated by CRISPR/Cas9-mediated genome editing. A synthetic *twk-40* crRNA, sgTwk-40 5′-CGATATCGGCAACAGTTATA-3′, was duplexed with tracrRNA and complexed with Cas9 protein. The ribonucleoprotein complex was injected together with the single-stranded DNA repair template oNZ158 5′-GAGGAAAATTGCTCTGCATCCTATACGCACTTTTCGGTGTCCCACTGATCACT ATAACTGTTGCCGATATCGGAAAGTTTTTATCTGAAAATATTGTTCAATT-3′. The repair template introduced the L159T substitution in TWK-40/T28A8.1a.

### Generation of the conditional *pyk-1* allele

The conditional *pyk-1* allele, *pyk-1(syb10871 syb11010)*, was designed in this study and generated by SunyBiotech using CRISPR/Cas9-mediated genome editing. An nls::loxP::sl2 cassette was inserted immediately upstream of the start codon of *pyk-1a* (F25H5.3a), and a loxP::wScarlet cassette was inserted immediately upstream of the stop codon, thereby flanking the *pyk-1a* coding region with loxP sites. Cre expression was used to induce conditional deletion of pyk-1. Loss of PYK-1::wScarlet fluorescence was assessed in AVA neurons.

The targeted *pyk-1a* isoform is the longest isoform of *pyk-1*. Cre-mediated recombination removes the intervening genomic sequence, including intronic regions that may contain regulatory elements required for PYK-1 expression in AVA. This could explain the loss of wScarlet fluorescence rather than its nuclear relocalisation following recombination.

### PYK-1 overexpression

Wild-type *pyk-1* and the corresponding mutant were expressed from extrachromosomal arrays using an approximately 10-kb genomic fragment encompassing the *pyk-1* genomic region and endogenous regulatory sequences. The amino-acid substitution was introduced by site-directed mutagenesis.

### Live fluorescence microscopy

#### Synaptic marker imaging

Live L4 larvae were immobilised on dry agarose pads in a small drop of M9 buffer. To assess potential genotype-dependent changes in promoter-driven fluorescence, we examined GFP fluorescence intensity during FACS analysis and in additional fluorescent reporter lines, including the RFP reference channel used for AVA calcium imaging^23^. No obvious genotype-dependent differences were observed in these assessments. Fluorescence signals were captured using a Nikon Eclipse 90i confocal microscope and Nikon Eclipse Ti2 with Yokogawa CSU-X1 spinning disc field scanning or Zeiss Airyscan scanning confocal system with 40× or 63× objectives. All fluorescent images were analysed by the in-house-developed puncta analyser^86^ or ImageJ intensity analyser. Fluorescence intensity was quantified as the mean pixel intensity within each identified punctum. The mean punctum intensity was then calculated for each animal, with individual animals used as independent biological replicates for statistical analysis.

### Immunofluorescent staining and imaging

Mixed-stage animals with endogenously tagged *twk-40* (*twk-40*::tagRFP::ZF) and the synaptic vesicle marker (P*twk-40-*SNB-1::GFP) were used to perform immunofluorescent staining as previously described^87^. Primary antibodies against tagRFP (Thermo Fisher Scientific, R10367, rabbit) and/or green fluorescent protein (GFP) (Roche, mouse) were used at 1:500 and 1:200 dilutions, respectively. Goat anti-rabbit (Alexa Fluor 594) and/or goat anti-mouse (Alexa Fluor 488) secondary antibodies were used at a 1:5000 dilution. Images were acquired with a Nikon spinning disc confocal microscope with a 63× objective and reconstructed by maximum intensity projection. Single layers of acquired images from animals co-expressing TWK-40::tagRFP::ZF and SNB-1::GFP were examined for subcellular colocalisation using an ImageJ plugin (JACoP).

### Spontaneous locomotion analyses

Freely moving animals were recorded as previously described^23^. Movies were analysed as previously described^23,28,88^. Briefly, at 10 fps, forward movement is defined as speed > 1 pixel/s, backward movement as speed < −1 pixel/s, and pausing as −1 ≤ speed ≤ 1 pixel/s. All animal data recorded on the same day were combined to perform the analysis.

### Heat shock induction of degradation

Transgenic *C. elegans* expressing *zif-1* under the control of the *hsp-16.2* promoter were heat shocked at 37°C for 30 minutes to induce targeted protein degradation. After heat shock, animals were returned to 20°C and allowed to develop until the L4 assay time point. Locomotor behaviour was manually quantified under a dissecting microscope over a 1-minute time window by scoring the duration of forward movement, backward movement, and pausing. The timing of the induced degradation paradigm was based on previously characterised ZIF-1/ZF1 kinetics, which showed rapid depletion of ZF1-tagged proteins within approximately 30–45 min following heat-shock-induced ZIF-1 expression.

### Behaviours upon optogenetic stimulation

An individual L4-stage animal was transferred to an imaging plate with a thin layer of evenly spread OP50. LED light stimulation was applied to activate Chrimson in AVA. Animals were cultured on NGM plates with or without ATR for at least 1 day for behaviour recording. During stimulation, L4-stage animals were exposed to 574-nm LED light (Thorlabs) at the maximum controller setting, with a 10-s pre-stimulation, 10-s stimulation, and 10-s post-stimulation cycle.

### Behaviours upon head touch stimulation

A platinum wire pick was used to gently touch the head of the animal to elicit a reversal response. Backward movement following stimulation was manually scored under a dissecting microscope. The experimenter was aware of the wild-type controls but was blinded to the identities of the different mutant genotypes during behavioural scoring.

### Simultaneous calcium recording and optogenetic stimulation

The experimental process was previously described^23,88^. Briefly, L4-stage animals carrying AVA calcium reporters and AVA-specific Chrimson were incubated on ATR(-) or ATR(+) plates for at least 1 day. They were immobilised on dry agarose pads in M9 buffer and subjected to full-spectrum light illumination for simultaneous calcium imaging and Chrimson-mediated activation at 10 fps or 20 fps for 6 to 20 s. GCaMP and RFP fluorescence intensities were measured from the AVA soma. Because the illumination used for calcium imaging also activated Chrimson, the GCaMP/RFP ratio at the first acquired frame (t = 0 s) was used as an operational baseline, assuming that the calcium response had not yet developed at this time point. ΔF/F was calculated relative to this baseline.

### Behavioural analyses upon chemogenetic manipulation

L4-stage animals carrying AVA-HisCl::mCherry^44^ were incubated on 0, 1, or 5 mM histamine plates with OP50 for 1 h. Backward movement in response to head touch was then scored as described above.

### Synaptic imaging upon chemogenetic manipulation

Animals carrying an integrated chromosomal array for AVA-HisCl::mCherry were cultured on 0, 1, or 5 mM histamine plates with OP50 for two generations. L4-stage animals from the second generation were used for synaptic imaging and quantification.

### Calcium imaging of premotor interneurons in moving animals

For AVA, an individual transgenic L3-stage animal expressing the GCaMP6s calcium sensor in AVA was mounted on a 2% NGM gel pad with ∼1 μL of M9 buffer so that it could move under a coverslip. Mounted animals were allowed to recover for 3 min on the pad before recording. The recording was performed using a 20× objective on a Zeiss inverted microscope equipped with an automated tracking system. Animals were automatically maintained within the field of view while the AVA soma was kept in focus. GCaMP and RFP fluorescence intensities were measured from the AVA soma, and the GCaMP/RFP ratio was calculated for each frame using in-house-developed MATLAB scripts^88^.

### Potassium imaging

Transgenic *C. elegans* expressing the FRET-based potassium sensor Kirin^49^ in AVA were immobilised on dry agarose pads in a small drop of M9 buffer. After a 5-minute recovery, ratiometric imaging was performed using a 20× objective on a Zeiss Axio Imager equipped with a dual-view optical splitter and a Hamamatsu camera. CFP was excited at ∼430 nm, and emission signals were collected simultaneously for CFP and YFP channels to compute the FRET ratio (YFP/CFP) to quantify intracellular K⁺ dynamics.

### RNA-seq and analysis

A transgenic *C. elegans* line expressing cytosolic GFP under the *twk-40* promoter (*hpIs860,* which is expressed in AVA, AVB, AVE, SAB, DVA, and anterior A-type motor neurons) was generated by integrating an extrachromosomal array and outcrossing. This line was crossed into *twk-40* loss-and gain-of-function backgrounds. Synchronised L4 larvae were dissociated using SDS-DTT and pronase treatment, followed by filtration and FACS to isolate GFP^+^ neurons. RNA was extracted using the Quick-RNA Microprep Kit, and libraries were prepared with SMART-Seq v4 and Nextera XT kits. Sequencing was performed on an Illumina NovaSeq platform. Reads were quality-checked (FastQC), aligned to the *C. elegans* genome (WBcel235) using STAR, and quantified with HTSeq. Differential expression analysis was performed using DESeq2, with genes considered significant at adjusted P < 0.05 and |log₂FC| ≥ 2 (Tables S1–S2). PCA and heatmaps were generated in R. Gene Ontology (GO) biological process overrepresentation analysis was performed using the PANTHER Overrepresentation Test (GO annotation release: 28 April 2026). The analysis included 776 differentially expressed genes from *twk-40(gf)* animals, of which 687 were mapped by PANTHER. Genes with non-missing adjusted P values in the RNA-seq differential expression analysis were used as the reference set (28,147 genes, of which 19,055 were mapped by PANTHER). Enrichment was assessed using Fisher’s exact test with false discovery rate (FDR) correction for multiple testing. GO terms with FDR < 0.05 were considered significant (Tables S3–S5).

### Auxin-induced protein degradation

Targeted protein degradation was performed using the auxin-inducible degradation system as previously described^59,60^. Animals carrying the corresponding AID-tagged transcription factors, with or without AVA-specific TIR1 expression, were transferred to NGM plates containing 4 mM 1-naphthaleneacetic acid potassium salt (K-NAA) at the L3 larval stage and incubated overnight at 20°C before synaptic imaging at L4.

### Statistical analyses

Statistical analyses were performed using GraphPad Prism 9.5.1. Normality was assessed using the Shapiro–Wilk test, and homogeneity of variances was assessed using the Brown–Forsythe test. Independent two-group comparisons were analysed using the two-tailed Mann–Whitney U test. Paired calcium measurements at illumination onset and 6 s later were analysed using the two-tailed Wilcoxon matched-pairs signed-rank test with Holm correction for multiple comparisons. For comparisons among multiple groups, ordinary one-way ANOVA was used when normality and equal-variance assumptions were satisfied, and Brown–Forsythe and Welch ANOVA were used when variances were unequal. Data that did not meet the normality assumption were analysed using the Kruskal–Wallis test. Post hoc multiple comparisons were performed using Dunn’s test following Kruskal–Wallis tests, Sidak’s test following ordinary one-way ANOVA, and Dunnett’s T3 test following Brown–Forsythe and Welch ANOVA. Fisher’s exact test was used for categorical behavioural responses. Statistical significance was defined as P < 0.05.

Brown–Forsythe and Welch ANOVA were used for the GCaMP/RFP ratio (Fig. 4C), Kirin FRET ratio (Fig. 5A), PYK-1 overexpression experiments (Fig. 6A), TWK-40 gain-of-function allele comparison (Fig. S4B), and UNC-42::GFP fluorescence intensity (Fig. 8B). Ordinary one-way ANOVA was used for FAX-1::EGFP fluorescence intensity (Fig. 8B). The remaining multiple-group fluorescence intensity comparisons were analysed using the Kruskal–Wallis test. Auxin-induced degradation experiments (Fig. 8D–F) were analysed using two-tailed Mann–Whitney U tests.

---

**Figure S1.**
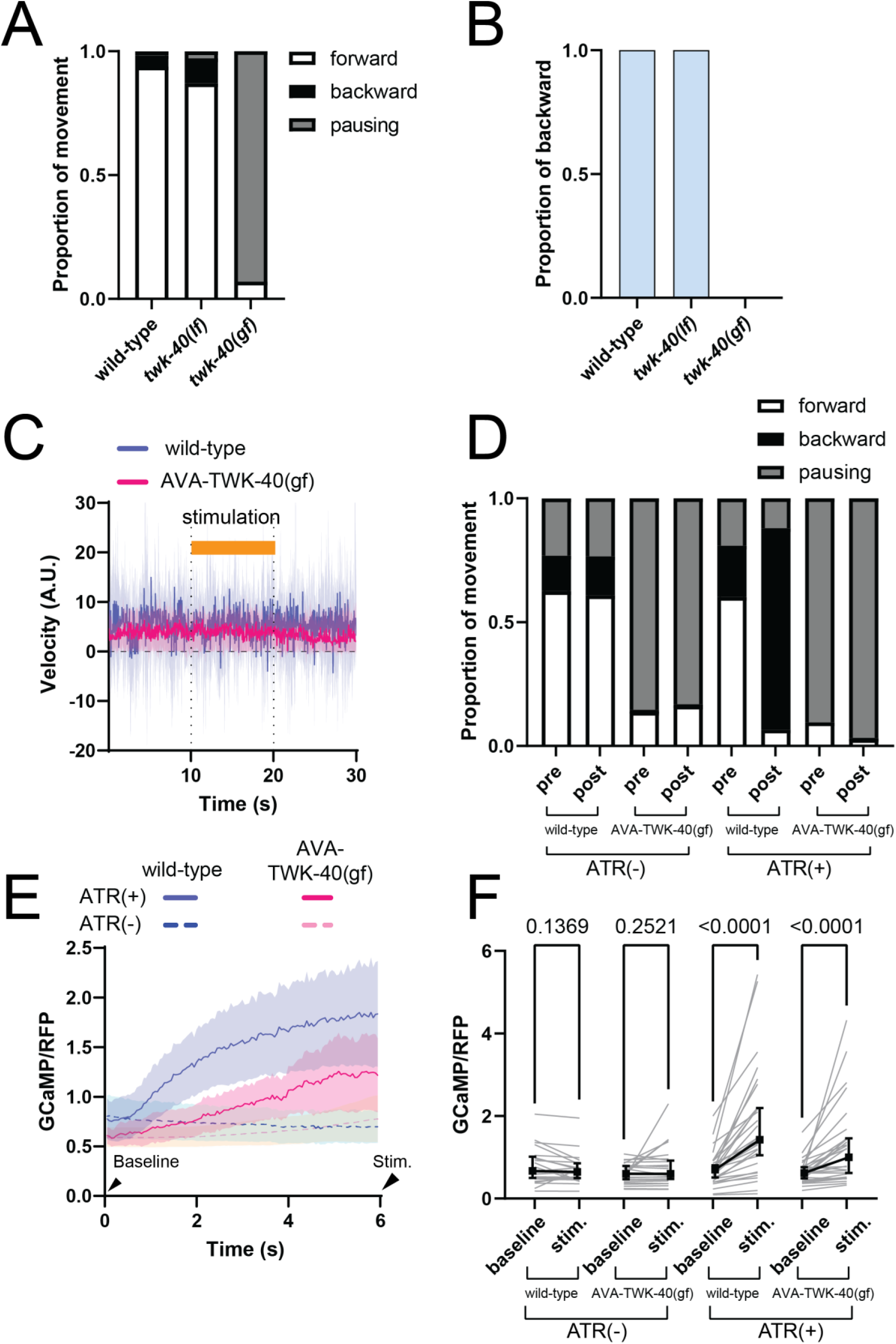
TWK-40 regulates AVA activity and locomotion. (A) Proportion of movement (forward, backward, pausing) during spontaneous locomotion for each genotype, pooled across 10 animals per genotype. Loss of twk-40 (twk-40(lf)) increases the proportion of backward movement, whereas gain of twk-40 (twk-40(gf)) abolishes backward movement and increases the pausing state. **Sample sizes:** N = 10 animals per genotype. (B) Proportion of animals exhibiting a backward escape response upon head touch for each genotype. Backward movement is blocked in twk-40(gf) mutants, indicating impaired reversal behaviour. Statistical comparisons were performed using Fisher’s exact test. Wild-type and twk-40(lf) animals did not differ in their backward response (P > 0.9999), whereas twk-40(gf) animals showed a significantly reduced backward response compared with wild-type (P < 0.0001). **Sample sizes:** N = 50 animals per genotype. (C) Velocity traces of each genotype without ATR supplementation, showing no difference in movement upon light stimulation. **Sample sizes:** wild-type (N = 12 animals), AVA-TWK-40(gf) (N = 18 animals). (D) Proportion of movement (forward, backward, pausing) for each condition in each genotype, pooled across the respective animals for each genotype. Animals without ATR supplementation show no changes in movement proportion within each genotype. Wild-type animals supplemented with ATR increase backward movement upon stimulation, whereas AVA-TWK-40(gf) animals exhibit an increased pausing state instead of backward movement. **Sample sizes:** ATR(-): wild-type (N = 12 animals), AVA-TWK-40(gf) (N = 18 animals). ATR(+): wild-type (N = 10 animals), AVA-TWK-40(gf) (N = 11 animals). (E) Calcium traces (GCaMP/RFP) over time in AVA neurons with ATR(+) or without ATR(-) supplementation. Stimulation begins at the onset of recording and continues throughout the recording period, as the imaging illumination simultaneously activates Chrimson. AVA stimulation in the presence of ATR increases calcium levels in both wild-type and AVA-TWK-40(gf) animals. However, the increase is attenuated in AVA-TWK-40(gf) mutants compared to wild-type. (E, F) Baseline indicates the first frame at stimulation onset; Stim. indicates a later time point during Chrimson stimulation. **Sample sizes:** ATR(-): wild-type (N = 22 animals), AVA-TWK-40(gf) (N = 25 animals). ATR(+): wild-type (N = 33 animals), AVA-TWK-40(gf) (N = 30 animals). (F) Quantification of GCaMP/RFP fluorescence at the beginning (0 s) and after 6 s of continuous illumination in (E). AVA stimulation significantly increases calcium levels in both wild-type and AVA-TWK-40(gf) animals when ATR is supplemented. AVA-TWK-40(gf) animals show a small but significant increase in calcium during illumination relative to the baseline measured at illumination onset. Two-tailed Wilcoxon matched-pairs signed-rank tests with Holm correction were used.

**Figure S2.**
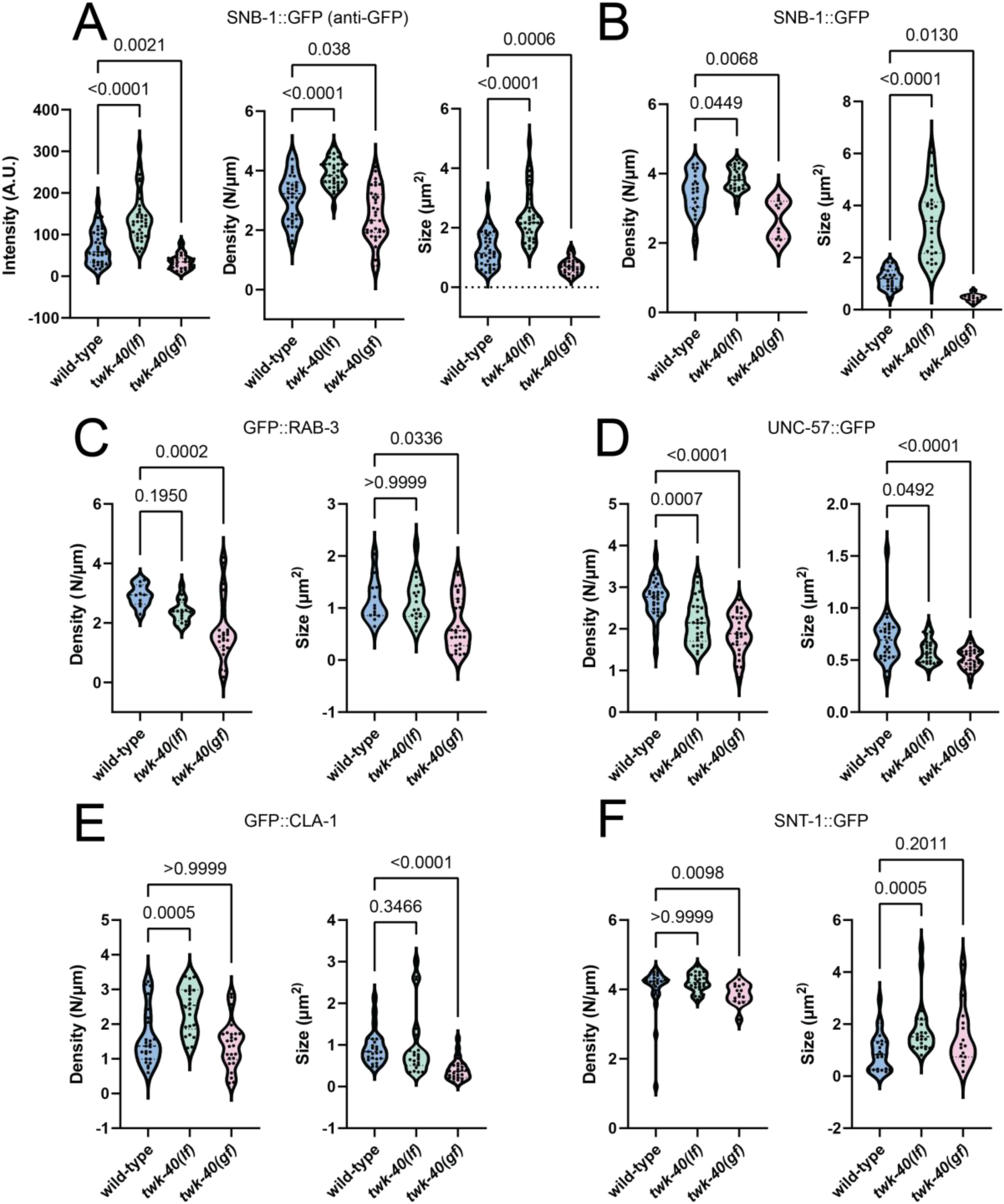
TWK-40 activity affects presynaptic organisation. (A) Quantification of anti-GFP immunostaining for AVA-specific SNB-1::GFP puncta intensity (Left), puncta density (Middle) and puncta size (Right) along the ventral nerve cord in wild-type, twk-40(lf), and twk-40(gf) animals. Each SNB-1::GFP parameter is significantly increased in twk-40(lf) mutants and decreased in twk-40(gf) mutants. **Sample sizes:** wild-type (N = 35 animals), twk-40(lf) (N = 36 animals), twk-40(gf) (N = 35 animals). (B–F) Quantification of puncta density (Left) and puncta size (Right) for SNB-1::GFP, GFP::RAB-3, UNC-57::GFP, GFP::CLA-1, and SNT-1::GFP along the ventral nerve cord in wild-type, twk-40(lf), and twk-40(gf) animals, showing marker-and parameter-specific changes in puncta density and size. **Sample sizes:** SNB-1::GFP: wild-type (N = 22 animals), twk-40(lf) (N = 25 animals), twk-40(gf) (N = 12 animals). GFP::RAB-3: wild-type (N = 10 animals), twk-40(lf) (N = 16 animals), twk-40(gf) (N = 24 animals). UNC-57::GFP: wild-type (N = 28 animals), twk-40(lf) (N = 26 animals), twk-40(gf) (N = 28 animals). GFP::CLA-1: wild-type (N = 25 animals), twk-40(lf) (N = 22 animals), twk-40(gf) (N = 16 animals). SNT-1::GFP: wild-type (N = 25 animals), twk-40(lf) (N = 24 animals), twk-40(gf) (N = 15 animals).

**Figure S4.**
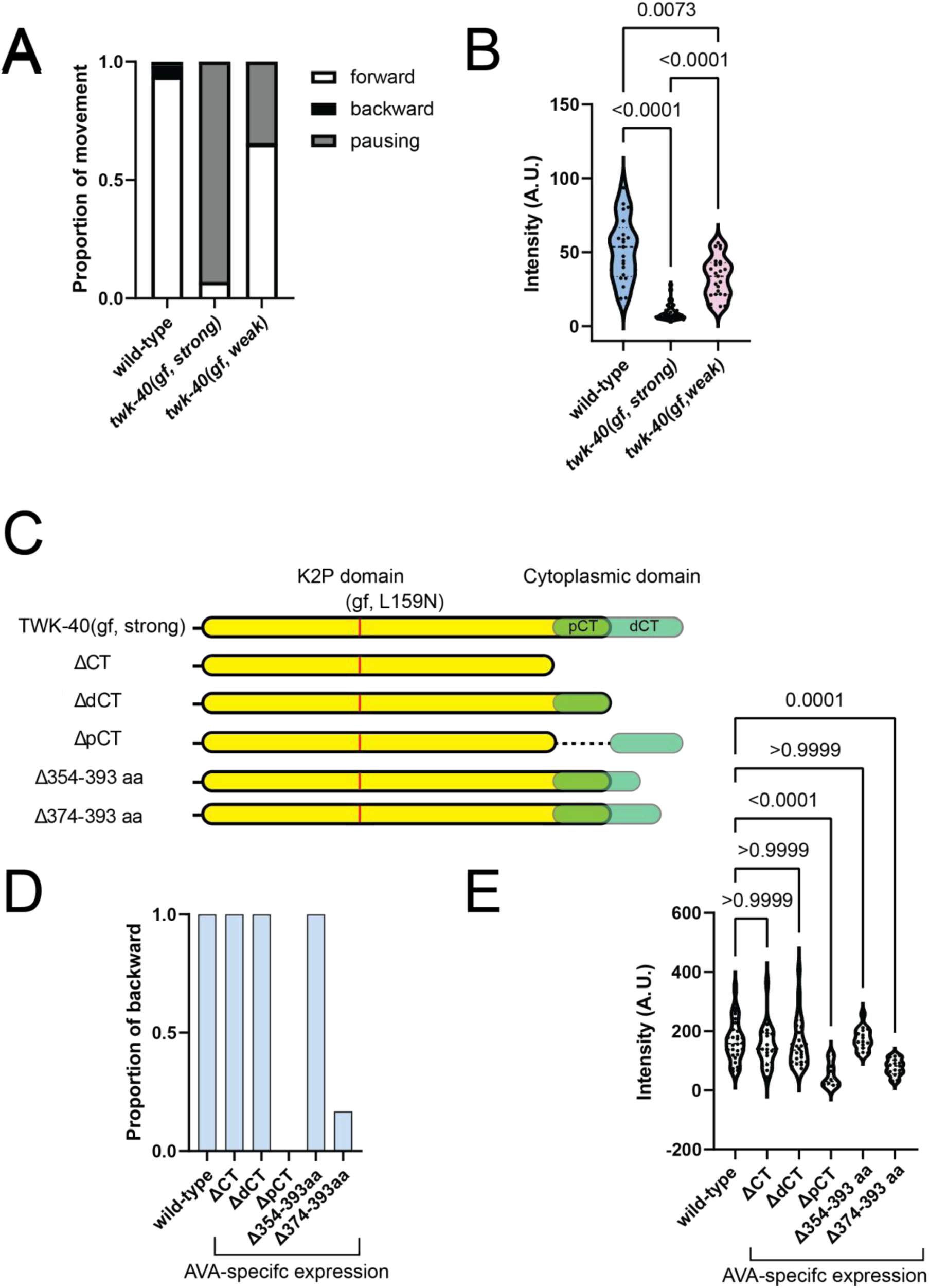
TWK-40 channel activity tracks presynaptic organisation. (A) Proportion of movement (forward, backward, pausing) for each genotype, showing that strong and weak alleles of twk-40(gf) exhibit distinct locomotor phenotypes, pooled across 10 animals per genotype. **Sample sizes:** N = 10 animals per genotype. (B) Quantification of AVA-SNB-1::GFP puncta intensity along the ventral nerve cord in each genotype. twk-40(gf, strong) exhibits a significant reduction in SNB-1::GFP intensity, while twk-40(gf, weak) shows a moderate but still significant reduction. **Sample sizes:** wild-type (N = 21 animals), twk-40(gf, strong) (N = 51 animals), twk-40(gf, weak) (N = 25 animals). (C) Schematic of TWK-40(gf, strong) C-terminal truncation constructs. For truncation analysis, the cytoplasmic C-terminal tail was operationally divided into a proximal C-terminal region (pCT) and a distal C-terminal region (dCT); these designations do not imply established structural subdomains. ΔCT removes the entire cytoplasmic tail, whereas ΔpCT and ΔdCT selectively remove the proximal or distal region. Smaller deletions within the distal C-terminal region were generated to further map sequences required for TWK-40-dependent presynaptic regulation. (D) Proportion of animals exhibiting a backward escape response upon head touch for each genotype, showing that backward escape responses are abolished in animals overexpressing TWK-40(gf, ΔpCT) and markedly reduced in animals expressing TWK-40(gf, Δ374–393 aa). Fisher’s exact test: wild-type vs ΔCT, P = 1; wild-type vs ΔdCT, P = 1; wild-type vs Δ374–393 aa, P < 0.0001; wild-type vs Δ354–393 aa, P = 1; wild-type vs ΔpCT, P < 0.0001. **Sample sizes:** N = 30 animals per genotype. (E) Quantification of AVA-SNB-1::GFP puncta intensity along the ventral nerve cord in each genotype. Overexpression of twk-40(gf, ΔpCT) and twk-40(gf, Δ374–393 aa) significantly reduces SNB-1::GFP intensity. **Sample sizes:** wild-type (N = 28 animals), ΔCT (N = 20 animals), ΔdCT (N = 28 animals), ΔpCT (N = 10 animals), Δ354–393 aa (N = 15 animals), Δ374–393 aa (N = 15 animals).

**Figure S7.**
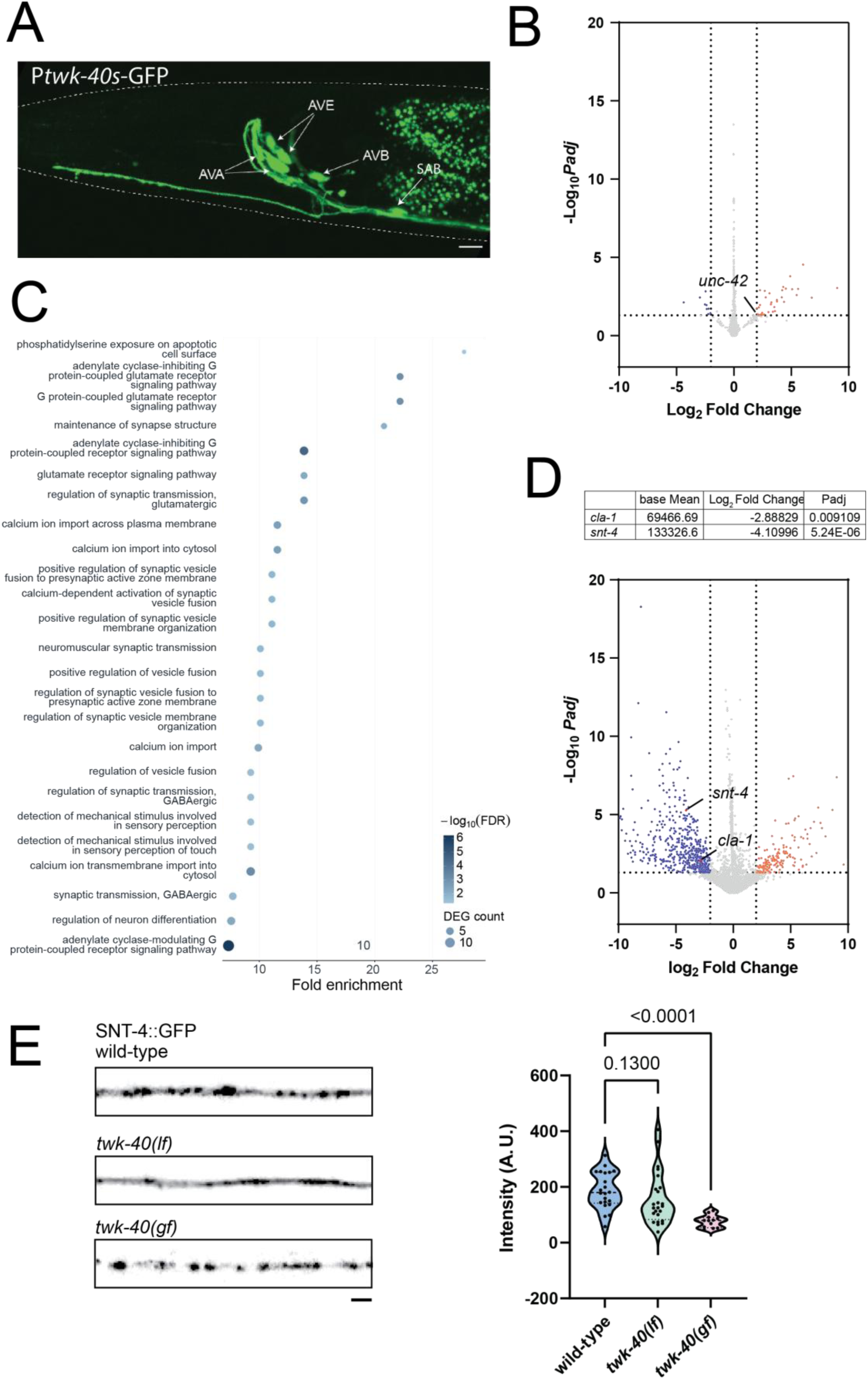
twk-40(gf) alters the neuronal transcriptome and reduces presynaptic protein levels. (A) Representative confocal image of Ptwk-40s::GFP expression, labelling a subset of neurons, including AVA, AVB, AVE, and SAB. Scale bar: 5 μm. (B) Volcano plot of differentially expressed genes in twk-40-expressing neurons from twk-40(lf) animals, showing fewer significant transcriptional changes than in twk-40(gf) (Table S2). **Sample size:** wild-type control (N = 3 biological replicates); twk-40(lf) (N = 3 biological replicates) (C) Gene Ontology (GO) biological process overrepresentation analysis of differentially expressed genes in twk-40(gf) animals using the same RNA-seq-derived reference set as in Figure 7C. The 25 significantly overrepresented GO terms with the highest fold enrichment are shown. The x-axis represents fold enrichment, dot size indicates the number of differentially expressed genes associated with each GO term, and colour represents −log₁₀(FDR) (Tables S4 and S5). (D) **Top:** Expression data for cla-1 and snt-4 displaying base mean expression, log2 fold change, and adjusted p-values. **Bottom:** Volcano plot of differentially expressed genes in twk-40-expressing neurons from twk-40(gf) animals. Notable downregulated genes include cla-1 and snt-4, which are associated with presynaptic function. padj = adjusted P value (E) **Left:** Representative confocal images of SNT-4::GFP fluorescence at AVA synapses in wild-type, twk-40(lf), and twk-40(gf) animals. Scale bar: 2 μm. **Right:** Quantification of SNT-4::GFP fluorescence intensity, showing a significant decrease in twk-40(gf). **Sample sizes:** wild-type (N = 22 animals), twk-40(lf) (N = 24 animals), twk-40(gf) (N = 12 animals).

## Notes

### Competing Interest Statement

The authors have declared no competing interest.

### Summary of Updates

We have revised the manuscript to improve clarity and consistency, updated several figures and their legends, clarified experimental controls and statistical analyses, and refined the Methods and supplementary materials. The main findings and conclusions remain unchanged.

