## Supplemental Table S6 for "K2P Channels Regulate Presynaptic Organisation through a Membrane Potential-Independent Mechanism"

Strains and Plasmids Used in This Study

| Strain | Allele | Genotype | Plasmid | Figures |
| --- | --- | --- | --- | --- |
| ZM10539 | *hpIs758;*  *hpIs774* | P*rig-3* LoxP EBFP stop LoxP Chrimson::RFP  *Ptwk-40s-*Cre;  *Ptwk-40s-*GCaMP6s::mNeptune | pJH4253  pJH4237  pJH4293 | Figure 1A, B; Figure S1 C-F |
| ZM10888 | *hpIs768;*  *hpIs758;*  *hpIs775* | P*flp-*18 LoxP EBFP stop LoxP TWK-40(L159N)::GFP  *Ptwk-40s-Cre;*  P*rig-3* LoxP EBFP stop LoxP Chrimson::RFP  P*twk-40s-*Cre;  P*twk-40s-*GCaMP6s::mNeptune | pJH4316  pJH4253  pJH4237  pJH4293 | Figure 1A, B;  Figure S1 C–F |
| ZM11287 | *hpEx4479* | *Pnpr-4-*SNB-1::pHluorin **lin-15* | pJH4783 | Figure 1C |
| ZM11284 | *twk-40(bln336);*  *hpEx4479* | *twk-40(gf, L159N)*;  P*npr-4-*SNB-1::pHluorin **lin-15* | pJH4783 | Figure 1C |
| ZM11285 | *twk-40(hp834);*  *hpEx4479* | *twk-40(lf);*  P*npr-4-*SNB-1::pHluorin **lin-15* | pJH4783 | Figure 1C |
| N2 | Wild-type | Wild-type | N/A | Figure S1 A, B; Figure 4B, Figure S4A; |
| ZM8302 | *twk-40(hp834)* | *twk-40(lf)* | N/A | Figure S1A, B |
| JIP1514 | *twk-40(bln336)* | *twk-40(gf, L159N)* | N/A | Figure S1A, B |
| AQ5556 | *twk-40(bln282);*  *hpIs833* | *twk-40*::tagRFP::ZF;  P*twk-40*-SNB-1::GFP **lin-15* | pJH4602-2 | Figure 2A |
| ZM10040 | *hpIs721* | P*rig-*3 FRT stop FRT SNB-1::GFP  P*nmr-1*-FLP  ***P*myo-2-*wCherry | pJH4169  pJH2673 | Figure 2B; Figure S2A, B; Figure S4B; Figure 4D; Figure 5B;  Figure 6A;  Figure 8C |
| ZM10417 | *twk-40(bln336);*  *hpIs721* | *twk-40(gf, L159N);*  P*rig-*3 FRT stop FRT SNB-1::GFP  P*nmr-1*-FLP  ***P*myo-2-*wCherry | pJH4169  pJH2673 | Figure 2B; Figure S2A, B; Figure S4B;  Figure 4D;  Figure 5D |
| ZM10416 | *twk-40(hp834);*  *hpIs721* | *twk-40(lf);* P*rig-*3 FRT stop FRT SNB-1::GFP  P*nmr-1*-FLP  ***P*myo-2-*wCherry | pJH4169  pJH2673 | Figure 2B; Figure S2A, B,  Figure 4D; Figure 5C; Figure 8C |
| ZM11258 | *twk-40(bln336 bln282); hpIs721;*  *hpEx3904* | *twk-40(gf, L159N)::ZF;* P*rig-*3 FRT stop FRT SNB-1::GFP  P*nmr-1*-FLP  ***P*myo-2-*wCherry;  P*rig-3 sl2* FRT EBFP FRT *zif-1;*  P*nmr-1-*FLP **HygR* | pJH4169  pJH2673  pJH4018 | Figure 2B |
| AQ5424 | *ljEx1749* | P*flp-*18 LoxP EBFP stop LoxP GFP::RAB-3  P*twk-40s-*Cre  **cc::mKate* | pCDB11  pJH4237 | Figure 2C; Figure S2C |
| AQ5425 | *twk-40(hp834); ljEx1749* | *twk-40(lf);* P*flp-*18 LoxP EBFP stop LoxP GFP::RAB-3  P*twk-40s-*Cre  **cc::mKate* | pCDB11  pJH4237 | Figure 2C; Figure S2C |
| AQ5426 | *twk-40(bln336); ljEx1749* | *twk-40(gf, L159N)*; P*flp-*18 LoxP EBFP stop LoxP GFP::RAB-3  P*twk-40s-*Cre  **cc::mKate* | pCDB11  pJH4237 | Figure 2C; Figure S2C |
| ZM11037 | *hpIs838* | P*npr-4*-UNC-57::GFP | pJH4623 | Figure 2D; Figure S2D |
| ZM11057 | *twk-40(hp834); hpIs838* | *twk-40(lf)*; P*npr-4*-UNC-57::GFP | pJH4623 | Figure 2D; Figure S2D |
| ZM11051 | *twk-40(bln336); hpIs838* | *twk-40(gf, L159N);* P*npr-4*-UNC-57::GFP | pJH4623 | Figure 2D; Figure S2D |
| AQ5427 | *ljEx1450* | P*flp-18* LoxP EBFP stop LoxP CLA-1::GFP  P*twk-40s-*Cre  **cc::mKate* | pCDB12  pJH4237 | Figure 2E; Figure S2E |
| AQ5428 | *twk-40(hp834); ljEx1450* | *twk-40(lf);* P*flp-18* LoxP EBFP stop LoxP CLA-1::GFP  P*twk-40s-*Cre  **cc::mKate* | pCDB12  pJH4237 | Figure 2E; Figure S2E |
| AQ5429 | *twk-40(bln336); ljEx1450* | *twk-40(gf, L159N)*; P*flp-18* LoxP EBFP stop LoxP CLA-1::GFP  P*twk-40s-*Cre  **cc::mKate* | pCDB12  pJH4237 | Figure 2E; Figure S2E |
| AQ5430 | *ljEx1451* | P*flp-18* LoxP EBFP stop LoxP SNT-1::GFP  P*twk-40s-*Cre  **HygR* | pCDB6  pJH4237 | Figure 2F; Figure S2F |
| AQ5431 | *twk-40(hp834);*  *ljEx1451* | *twk-40(lf)*; P*flp-18* LoxP EBFP stop LoxP SNT-1::GFP  P*twk-40s-*Cre  **HygR* | pCDB6  pJH4237 | Figure 2F; Figure S2F |
| AQ5432 | *twk-40(bln336); ljEx1451* | *twk-40(gf, L159N)*; P*flp-18* LoxP EBFP stop LoxP SNT-1::GFP  P*twk-40s-*Cre  **HygR* | pCDB6  pJH4237 | Figure 2F; Figure S2F |
| AQ5433 | *twk-40(bln336 bln282); hpIs721; ljEx1452* | *twk-40(gf, L159N)*::ZF; P*rig-3* FRT stop FRT SNB-1::GFP  P*nmr-1*-FLP ***P*myo-2-*wCherry;  P*hsp*-ZIF-1::RFP | pJH4169  pJH2673  pCDB13 | Figure 3B, C |
| ZM1079 | *nca-1 (hp102)* | *nca-1(gf)* | N/A | Figure 4B |
| ZM9955 | *twk-40(bln336 bln282)* | *twk-40(gf, L159N)*::ZF | N/A | Figure 4B |
| ZM10045 | *twk-40(bln336 bln282); nca-1(hp102)* | *twk-40(gf, L159N)*::ZF; *nca-1 (gf)* | N/A | Figure 4B |
| ZM9059 | *hpIs580* | P*rig-3*-GCaMP6s::wCherry **lin-15* | pJH3631 | Figure 4C |
| ZM8970 | *nca-1(hp102);*  *hpIs580* | *nca-1(gf);*  Prig-3-GCaMP6s::wCherry **lin-15* | pJH3631 | Figure 4C |
| ZM9818 | *twk-40(bln336); hpIs580* | *Twk-40(gf, L159N);* Prig-3-GCaMP6s::wCherry **lin-15* | pJH3631 | Figure 4C |
| ZM10064 | *twk-40(bln336 bln282); nca-1 (hp102);*  *hpIs580* | *twk-40(gf, L159N)*::ZF; *nca-1(gf);*  Prig-3-GCaMP6s::wCherry **lin-15* | pJH3631 | Figure 4C |
| ZM10415 | *nca-1(hp102); hpIs721* | *nca-1 (gf)*; P*rig-3* FRT stop FRT SNB-1::GFP  P*nmr-1*-FLP ***P*myo-2-*wCherry | pJH4169  pJH2673 | Figure 4D |
| ZM10414 | *nca-2(gk5);*  *nca-1(gk9);*  *hpIs721* | *nca-2 (lf); nca-1 (lf);* P*rig-3* FRT stop FRT SNB-1::GFP  P*nmr-1*-FLP ***P*myo-2-*wCherry | pJH4169  pJH2673 | Figure 4D |
| ZM10106 | *twk-40(bln336 bln282);*  *nca-1(hp102); hpIs721* | *twk-40(gf, L159N)*::ZF*; nca-1(gf)*; P*rig-3* FRT stop FRT SNB-1::GFP  P*nmr-1*-FLP ***P*myo-2-*wCherry | pJH4169  pJH2673 | Figure 4D |
| ZM10894 | *nca-2(gk5) twk-40(hp834);*  *nca-1(gk9);*  *hpIs721* | *nca-2 (lf) twk-40(lf)*; *nca-1(lf)*; P*rig-3* FRT stop FRT SNB-1::GFP  P*nmr-1*-FLP ***P*myo-2-*wCherry | pJH4169  pJH2673 | Figure 4D |
| ZM11247 | *hpIs636;*  *hpIs721* | P*rig-3*-HisCl1::SL2::mCherry;  P*rig-3* FRT stop FRT SNB-1::GFP  P*nmr-1*-FLP ***P*myo-2-*wCherry | pJH4169  pJH2673 | Figure 4E, F |
| JIP1588 | *twk-40(bln404)* | *twk-40(gf, weak, L159T)* | N/A | Figure S4A |
| ZM10991 | *twk-40(bln404);*  *hpIs721* | *twk-40(gf, weak, L159T)*; P*rig-3* FRT stop FRT SNB-1::GFP  P*nmr-1*-FLP ***P*myo-2-*wCherry | pJH4169  pJH2673 | Figure S4B |
| AQ5434 | *hpIs721;*  *ljEx1453* | P*rig-3* FRT stop FRT SNB-1::GFP  P*nmr-1*-FLP ***P*myo-2-*wCherry; P*flp-18* LoxP EBFP stop LoxP TWK-40(L159N, ΔCT)  P*twk-40s-*Cre  ***P*myo-3-*mCherry | pJH4169  pJH2673  pCDB15  pJH4237 | Figure S4D, E |
| AQ5435 | *hpIs721; ljEx1454* | P*rig-3* FRT stop FRT SNB-1::GFP  P*nmr-1*-FLP ***P*myo-2-*wCherry; P*flp-18* LoxP EBFP stop LoxP TWK-40(L159N, ΔdCT)  P*twk-40s-*Cre  ***P*myo-3-*mCherry | pJH4169  pJH2673  pCDB16  pJH4237 | Figure S4D, E |
| AQ5436 | *hpIs721; ljEx1455* | P*rig-3* FRT stop FRT SNB-1::GFP  P*nmr-1*-FLP ***P*myo-2-*wCherry; P*flp-18* LoxP EBFP stop LoxP TWK-40(L159N, ΔpCT)  P*twk-40s-*Cre  **HygR* | pJH4169  pJH2673  pCDB17  pJH4237 | Figure S4D, E |
| AQ5437 | *hpIs721; ljEx1456* | P*rig-3* FRT stop FRT SNB-1::GFP  P*nmr-1*-FLP ***P*myo-2-*wCherry; P*flp-18* LoxP EBFP stop LoxP TWK-40(L159N, Δ354-393 aa)  P*twk-40s-*Cre  **HygR* | pJH4169  pJH2673  pCDB18  pJH4237 | Figure S4D, E |
| AQ5438 | *hpIs721; ljEx1457* | P*rig-3* FRT stop FRT SNB-1::GFP  P*nmr-1*-FLP ***P*myo-2-*wCherry; P*flp-18* LoxP EBFP stop LoxP TWK-40(L159N, Δ374-393 aa)  P*twk-40s-*Cre  **HygR* | pJH4169  pJH2673  pCDB19  pJH4237 | Figure S4D, E |
| AQ5439 | *ljEx1458* | P*flp-18* LoxP EBFP stop LoxP KIRIN  P*twk-40s-*Cre  **HygR* | pCDB20  pJH4237 | Figure 5A |
| AQ5440 | *twk-40(hp834); ljEx1458* | *twk-40(lf);* P*flp-18* LoxP EBFP stop LoxP KIRIN  P*twk-40s-*Cre  **HygR* | pCDB20  pJH4237 | Figure 5A |
| AQ5441 | *twk-40(bln336); ljEx1458* | *twk-40(gf, L159N);* P*flp-18* LoxP EBFP stop LoxP KIRIN  P*twk-40s-*Cre  **HygR* | pCDB20  pJH4237 | Figure 5A |
| AQ5442 | *eat-6(ad601); hpIs721* | *eat-6(lf);* P*rig-3* FRT stop FRT SNB-1::GFP  P*nmr-1*-FLP ***P*myo-2-*wCherry | pJH4169  pJH2673 | Figure 5B |
| AQ5443 | *kcc-2(ok3074); hpIs721* | *kcc-2(lf);* P*rig-3* FRT stop FRT SNB-1::GFP  P*nmr-1*-FLP ***P*myo-2-*wCherry | pJH4169  pJH2673 | Figure 5B |
| AQ5444 | *kcc-2(vs132); hpIs721* | *kcc-2(lf);* P*rig-3* FRT stop FRT SNB-1::GFP  P*nmr-1*-FLP ***P*myo-2-*wCherry | pJH4169  pJH2673 | Figure 5B |
| AQ5445 | *twk-40(hp834);*  *eat-6(ad601); hpIs721* | *twk-40(lf); eat-6(lf);* P*rig-3* FRT stop FRT SNB-1::GFP  P*nmr-1*-FLP ***P*myo-2-*wCherry | pJH4169  pJH2673 | Figure 5C |
| AQ5446 | *twk-40(hp834); kcc-2(ok3074); hpIs721* | *twk-40(lf); kcc-2(lf);* P*rig-3* FRT stop FRT SNB-1::GFP  P*nmr-1*-FLP ***P*myo-2-*wCherry; | pJH4169  pJH2673 | Figure 5C |
| AQ5447 | *twk-40(bln336); eat-6(ad601); hpIs721* | *twk-40(gf, L159N); eat-6(lf);* P*rig-3* FRT stop FRT SNB-1::GFP  P*nmr-1*-FLP ***P*myo-2-*wCherry | pJH4169  pJH2673 | Figure 5D |
| AQ5448 | *twk-40(bln336); kcc-2(ok3074); hpIs721* | *twk-40(gf, L159N); kcc-2(lf);* P*rig-3* FRT stop FRT SNB-1::GFP  P*nmr-1*-FLP ***P*myo-2-*wCherry | pJH4169  pJH2673 | Figure 5D |
| AQ5557 | *hpIs721; ljEx1801* | P*rig-3* FRT stop FRT SNB-1::GFP  P*nmr-1*-FLP ***P*myo-2-*wCherry; P*pyk-1L*-PYK-1 (gDNA)  **HygR* | pJH4169  pJH2673  pCDB21 | Figure 6A |
| AQ5558 | *hpIs721; ljEx1802* | P*rig-3* FRT stop FRT SNB-1::GFP  P*nmr-1*-FLP ***P*myo-2-*wCherry; P*pyk-1L*-PYK-1 (T121L)  **HygR* | pJH4169  pJH2673  pCDB22 | Figure 6A |
| AQ5559 | *pyk-1(syb10871 syb11010)* | nls::loxP::sl2::*pyk-1a*::loxP::wScarlet::stop | N/A | Figure 6C |
| AQ5560 | *pyk-1(syb10871 syb11010); hpEx4555* | nls::loxP::sl2::*pyk-1a*::loxP::wScarlet::stop; *Pnpr-4*-mito::GFP | pJH4934 | Figure 6C |
| AQ5561 | *pyk-1(syb10871 syb11010); ljEx1450* | nls::loxP::sl2::*pyk-1a*::loxP::wScarlet::stop; P*flp-18* LoxP EBFP stop LoxP CLA-1::GFP  P*twk-40s-*Cre  **cc::mKate* | pCDB12  pJH4237 | Figure 6C |
| AQ5562 | *pyk-1(syb10871 syb11010); hpIs721* | nls::loxP::sl2::*pyk-1a*::loxP::wScarlet::stop; P*rig-3* FRT stop FRT SNB-1::GFP  P*nmr-1*-FLP ***P*myo-2-*wCherry | pJH4169  pJH2673 | Figure 6D |
| AQ5563 | *pyk-1(syb10871 syb11010); hpIs721; ljEx1083* | nls::loxP::sl2::*pyk-1a*::loxP::wScarlet::stop; P*rig-3* FRT stop FRT SNB-1::GFP  P*nmr-1*-FLP ***P*myo-2-*wCherry;  P*flp-18-*Cre  **HygR* | pJH4169  pJH2673  pCDB23 | Figure 6D |
| AQ5564 | *pyk-1(syb10871 syb11010); twk-40(hp834); hpIs721* | nls::loxP::sl2::*pyk-1a*::loxP::wScarlet::stop; *twk-40(lf);* P*rig-3* FRT stop FRT SNB-1::GFP  P*nmr-1*-FLP ***P*myo-2-*wCherry | pJH4169  pJH2673 | Figure 6D |
| AQ5565 | *pyk-1(syb10871 syb11010); twk-40(hp834); hpIs721; ljEx1084* | nls::loxP::sl2::*pyk-1a*::loxP::wScarlet::stop; *twk-40(lf);* P*rig-3* FRT stop FRT SNB-1::GFP  P*nmr-1*-FLP ***P*myo-2-*wCherry; P*flp-18-*Cre  **HygR* | pJH4169  pJH2673  pCDB23 | Figure 6D |
| ZM11177 | *hpIs860* | P*twk-40s-*GFP | pJH4693 | Figure 7A, B; Figure S7A, C, D |
| ZM11208 | *twk-40(hp834); hpIs860* | *twk-40(lf);* P*twk-40s-*GFP | pJH4693 | Figure S7B |
| ZM11207 | *twk-40(bln336); hpIs860* | *twk-40(gf, L159N);* P*twk-40s-*GFP | pJH4693 | Figure 7 |
| AQ5455 | *ljEx1459* | P*flp-18* LoxP EBFP stop LoxP SNT-4::GFP  P*twk-40s-*Cre | pCDB8  pJH4237 | Figure S7E |
| AQ5456 | *twk-40(hp834);ljEx1459* | *twk-40(lf);* P*flp-18* LoxP EBFP stop LoxP SNT-4::GFP  P*twk-40s-*Cre | pCDB8  pJH4237 | Figure S7E |
| AQ5457 | *twk-40(bln336); ljEx1459* | *twk-40(gf, L159N);* P*flp-18* LoxP EBFP stop LoxP SNT-4::GFP  P*twk-40s-*Cre | pCDB8  pJH4237 | Figure S7E |
| AQ5458 | *unc-3(st12259)* | *unc-3::TY1::EGFP::3xFLAG* | N/A | Figure 8A, B |
| AQ5459 | *twk-40(hp834); unc-3(st12259)* | *twk-40(lf); unc-3::TY1::EGFP::3xFLAG* | N/A | Figure 8A, B |
| AQ5460 | *twk-40(bln336); unc-3(st12259)* | *twk-40(gf, L159N); unc-3::TY1::EGFP::3xFLAG* | N/A | Figure 8A, B |
| AQ5461 | *unc-42 (ot986)* | *unc-42::GFP* | N/A | Figure 8A, B |
| AQ5462 | *twk-40(hp834); unc-42 (ot986)* | *twk-40(lf); unc-42::GFP* | N/A | Figure 8A, B |
| AQ5463 | *twk-40(bln336); unc-42 (ot986)* | *twk-40(gf, L159N); unc-42::GFP* | N/A | Figure 8A, B |
| AQ5464 | *fax-1(wgIs164)* | *fax-1*::TY1::EGFP::3xFLAG **unc-119(+)* | N/A | Figure 8A, B |
| AQ5465 | *twk-40(hp834); fax-1(wgIs164)* | *twk-40(lf); fax-1*::TY1::EGFP::3xFLAG  **unc-119(+)* | N/A | Figure 8A, B |
| AQ5466 | *twk-40(bln336); fax-1(wgIs164)* | *twk-40(gf, L159N); fax-1*::TY1::EGFP::3xFLAG  **unc-119(+)* | N/A | Figure 8A, B |
| AQ5467 | *unc-3(e151); hpIs721* | *unc-3(lf); Prig-3 FRT stop FRT SNB-1::GFP*  *Pnmr-1-FLP *Pmyo-2-wCherry* | pJH4169  pJH2673 | Figure 8C |
| AQ5468 | *twk-40(hp834); unc-3 (e151); hpIs721* | *twk-40(lf); unc-3(lf);* P*rig-3* FRT stop FRT SNB-1::GFP  P*nmr-1*-FLP ***P*myo-2-*wCherry | pJH4169  pJH2673 | Figure 8C |
| AQ5469 | *unc-42 (e419); hpIs721* | *unc-42(lf);* P*rig-3* FRT stop FRT SNB-1::GFP  P*nmr-1*-FLP ***P*myo-2-*wCherry | pJH4169  pJH2673 | Figure 8C |
| AQ5470 | *twk-40(hp834); unc-42(e419); hpIs721* | *twk-40(lf); unc-42(lf);* P*rig-3* FRT stop FRT SNB-1::GFP;  P*nmr-1*-FLP*; **P*myo-2-*wCherry | pJH4169  pJH2673 | Figure 8C |
| AQ5471 | *fax-1(gm83); hpIs721* | *fax-1(lf);* P*rig-3* FRT stop FRT SNB-1::GFP  P*nmr-1*-FLP ***P*myo-2-*wCherry | pJH4169  pJH2673 | Figure 8C |
| AQ5472 | *twk-40(hp834); fax-1(gm83); hpIs721* | *twk-40(lf); fax-1(lf);* P*rig-3* FRT stop FRT SNB-1::GFP  P*nmr-1*-FLP ***P*myo-2-*wCherry | pJH4169  pJH2673 | Figure 8C |
| AQ5473  non-transgenic siblings | *twk-40(hp834); unc-3(ot837); hpIs721* | *twk-40(lf); unc-3::mNeonGreen::AID;*  P*rig-3* FRT stop FRT SNB-1::GFP  P*nmr-1*-FLP ***P*myo-2-*wCherry; | pJH4169  pJH2673 | Figure 8D |
| AQ5473 | *twk-40(hp834); unc-3(ot837); hpIs721; ljEx1460* | *twk-40(lf); unc-3::mNeonGreen::AID;*  P*rig-3* FRT stop FRT SNB-1::GFP  P*nmr-1*-FLP ***P*myo-2-*wCherry; P*rig-*3 FRT stop FRT TIR1::mRuby  **HygR* | pJH4169  pJH2673  pJH4190 | Figure 8D |
| AQ5474  non-transgenic siblings | *twk-40(hp834); unc-42(lg225); hpIs721* | *twk-40(lf); unc-42::AID;* P*rig-3* FRT stop FRT SNB-1::GFP  P*nmr-1*-FLP ***P*myo-2-*wCherry | pJH4169  pJH2673 | Figure 8E |
| AQ5474 | *twk-40(hp834); unc-42(lg225); hpIs721; ljEx1463* | *twk-40(lf); unc-42::AID;* P*rig-3* FRT stop FRT SNB-1::GFP  P*nmr-1*-FLP ***P*myo-2-*wCherry; *Prig-3 FRT stop FRT TIR1::mRuby*  **HygR* | pJH4169  pJH2673  pJH4190 | Figure 8E |
| AQ5475  non-transgenic siblings | *twk-40(hp834); fax-1(syb10141); hpIs721* | *twk-40(lf); AID::fax-1; Prig-3 FRT stop FRT SNB-1::GFP*  *Pnmr-1-FLP *Pmyo-2-wCherry* | pJH4169  pJH2673 | Figure 8F |
| AQ5475 | *twk-40(hp834); fax-1(syb10141); hpIs721; ljEx1462* | *twk-40(lf); AID::fax-1; Prig-3 FRT stop FRT SNB-1::GFP*  *Pnmr-1-FLP *Pmyo-2-wCherry*  P*rig-*3 FRT stop FRT TIR1::mRuby **HygR* | pJH4169  pJH2673  pJH4190 | Figure 8F |

* denotes coinjection markers.
